# The splicing factor hnRNP M is a critical regulator of innate immune gene expression in macrophages

**DOI:** 10.1101/617043

**Authors:** K.O. West, H.M. Scott, S. Torres-Odio, A.P. West, K.L. Patrick, R.O. Watson

## Abstract

While transcriptional control mechanisms of innate immune gene expression are well characterized, almost nothing is known about how pre-mRNA splicing decisions influence, or are influenced by, macrophage activation. Here, we demonstrate that the splicing factor hnRNP M is a critical repressor of innate immune gene expression and that its function is regulated by pathogen sensing cascades. Loss of hnRNP M leads to hyperinduction of a unique regulon of inflammatory and antimicrobial genes, including *IL6, Mx1*, and *Gbp5*, following a variety of innate immune stimuli. While mutating specific serines on hnRNP M had little effect on its ability to control pre-mRNA splicing or transcript levels of “housekeeping” genes in resting macrophages, it greatly impacted the protein’s ability to dampen induction of specific innate immune transcripts following activation of pathogen sensing cascades. These data reveal a previously unappreciated role for pattern recognition receptor signaling in controlling splicing factor phosphorylation and establish pre-mRNA splicing as a critical regulatory node in defining innate immune outcomes.

---

When innate immune cells like macrophages sense pathogens, they undergo a massive reprogramming of gene-expression. Decades of research have described the transcription factors and signal transduction cascades that initiate transcription of innate immune defense molecules. However, the contribution of post-transcriptional regulatory events and pre-mRNA splicing decisions to innate immune outcomes during macrophage activation remains understudied.

Multiple lines of evidence support a crucial role for pre-mRNA splicing regulation in determining innate immune gene expression outcomes. When primary mouse macrophages are treated with a Toll-like receptor 4 (TLR4) agonist to activate innate immune gene expression, individual transcripts show significant variation in the time it takes for them to be fully spliced, with some pre-mRNAs remaining unprocessed for hours after transcriptional activation^1,2^. Likewise, computational analyses of human primary macrophages reveal a robust increase in mRNA isoform diversity and a global preference for exon inclusion following lipopolysaccharide (LPS) treatment or *Salmonella enterica* serovar Typhimurium infection^3^. The production of functionally diverse protein isoforms via alternative splicing is also known to influence innate immune responses. Several important innate immune molecules downstream of pattern recognition receptors, such as the TLR adapter protein MyD88^4^, the interleukin-1 receptor associated kinase 1, IRAK1^5^, and even some of the TLRs themselves (TLR3, TLR4 co-receptor MD2)^6,7^, are regulated through expression of truncated isoforms that auto-inhibit full length protein function and dampen inflammatory responses. In the case of MyD88, splicing factors like SF3a1 have been directly implicated in generating the MyD88 short isoform (MyD88-S), which inhibits expression of pro-inflammatory cytokines like interleukin-6 (IL-6) following LPS treatment^8,9^. Despite these and other lines of evidence pointing to an important role for splicing and alternative splicing in controlling innate immune outcomes, little is known about how splicing decisions are regulated in innate immune cells undergoing gene expression reprogramming.

To date, a handful of RNA binding proteins (RBPs) have a characterized role in controlling innate immune gene expression, and in some cases, innate immune signaling has been connected to RBP function. For example, TLR4 signaling via LPS treatment promotes the shuttling of hnRNP U (heterogeneous nuclear ribonucleoprotein particle U) from the nucleus to the cytosol, resulting in differential expression of several innate immune cytokines (*Tnfα, IL6*, and *IL-1β*) via hnRNP U-dependent stabilization of cytosolic mRNAs^10^. Tristetraprolin (TTP), human antigen R (HUR), T-cell intracellular antigen 1 related protein (TIAR), and hnRNP K have also been implicated in controlling gene expression in LPS-activated macrophages, with TTP and HUR regulating mRNA decay and TIAR and hnRNP K causing translational repression^11–13^. Phosphorylation is generally thought to control subcellular localization and protein-protein interactions between these RBPs and others in the hnRNP and SR (serine-arginine rich) families^14–19^, but the kinases/phosphatases responsible for modifying them and the conditions under which these modifications are controlled remain poorly understood.

Two recent publications report changes to macrophage protein phosphorylation following infection with the intracellular pathogens bacterium *Mycobacterium tuberculosis*^20^ and *Cryptococcus neoformans*^21^. Intriguingly, a number of these differentially phosphorylated peptides were derived from splicing factors. In fact, “spliceosome” was the top over-represented phosphorylated pathway in *C. neoformans*-infected cells, suggesting that post-translational modification of splicing factors is critical for controlling innate immune responses to pathogens. One of the proteins that was significantly differentially phosphorylated in each of these datasets was hnRNP M. HnRNP M is a splicing factor and RBP that has been repeatedly implicated in cancer metastasis^22–25^ and muscle differentiation^26^. It has also been identified as a component of a large splicing regulatory complex containing Rbfox which controls alternative splicing in the brain^27^. Its role in regulating innate immune gene expression in macrophages is unknown, although interestingly, it is targeted by proteases from two positive strand RNA viruses (polio and coxsackievirus) in order to promote infection^28,29^. HnRNP M has also been found to influence dengue virus replication^30^, suggesting a role in regulating host antiviral responses.

Here, we demonstrate that abrogating hnRNP M expression in a macrophage cell line leads to hyperinduction of hundreds of transcripts following distinct innate immune stimuli, including infection with the gram-negative bacteria *Salmonella enterica* serovar Typhimurium, treatment with TLR2 and TLR4 agonists, and transfection of cytosolic dsDNA. While our data reveal that hnRNP M co-transcriptionally represses gene expression by influencing both constitutive and alternative splicing decisions, regulation of hnRNP M’s function via phosphorylation at S574 specifically controls the protein’s ability to inhibit intron removal of innate immune-activated transcripts. Consistent with its role in down-regulating macrophage activation, macrophages lacking hnRNP M were better able to control viral replication, emphasizing the importance of pre-mRNA splicing regulation in modulating the innate immune response to infection.

## RESULTS

### RNA-SEQ analysis reveals immune response genes are regulated by hnRNP M during *Salmonella* infection

To investigate a role for hnRNP M in regulating the innate immune response, we first tested how loss of hnRNP M globally influenced macrophage gene expression. Stable hnRNP M knockdown cell lines (hnRNP M KD) were generated by transducing RAW 264.7 mouse macrophages with lentiviral shRNA constructs designed to target hnRNP M or a control scramble (SCR) shRNA. Western blot and RT-qPCR analysis confirmed ~80% and 60% knockdown of hnRNP M using two different shRNA constructs (in KD1 and KD2, respectively) (Fig. 1a). Numerous attempts to knockout hnRNP M in RAW 264.7 macrophages by CRISPR/Cas9 gRNAs resulted exclusively in clones with in-frame insertions or deletions (data not shown). Therefore, we concluded that hnRNP M is essential in macrophages and continued our experiments using the viable knockdown cell lines.

**Figure 1.**
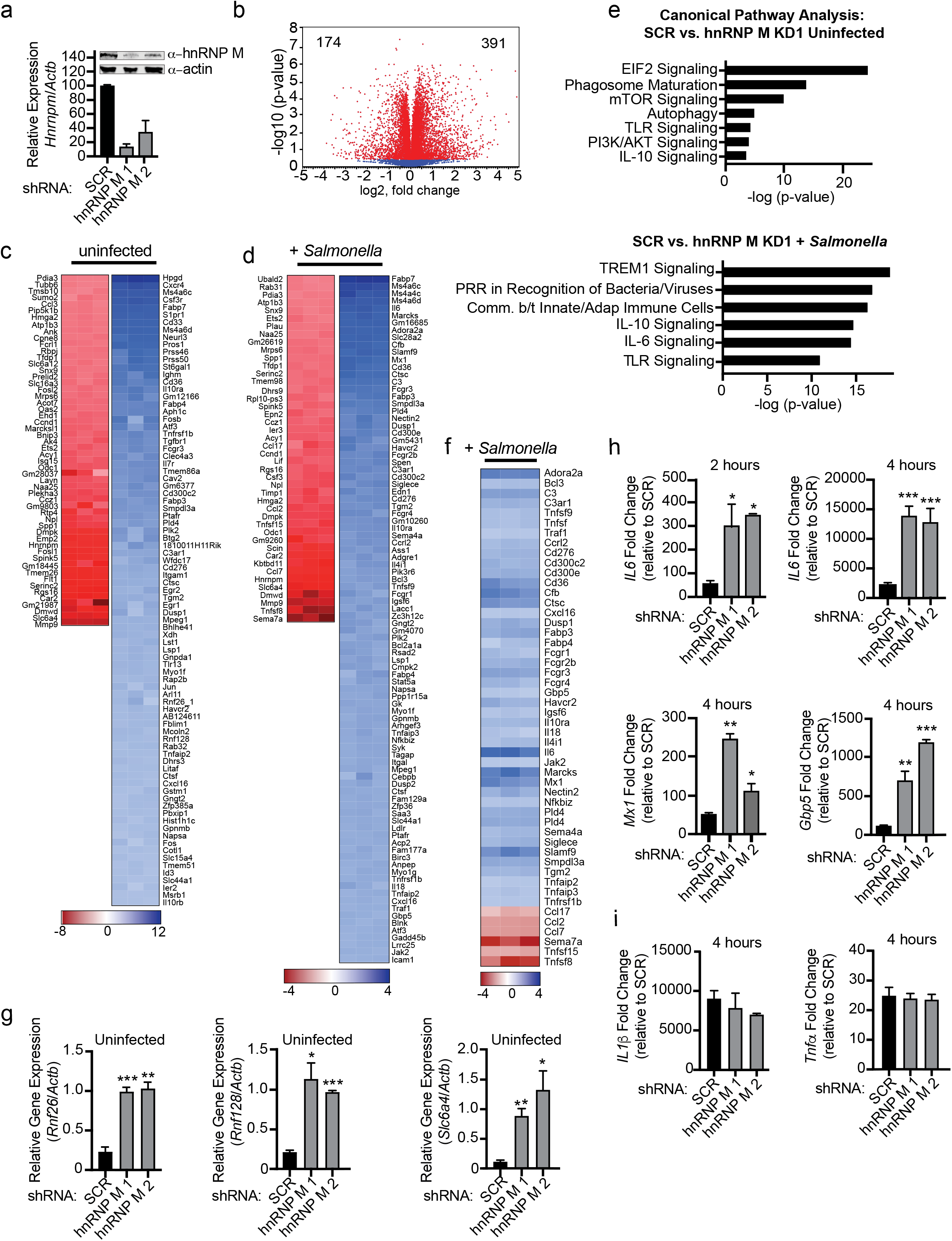
HnRNP M regulates expression of innate immune genes during *Salmonella* Typhimurium Infection. (a) Western analysis and RT-qPCR demonstrating effective depletion of hnRNP M in RAW 264.7 macrophages. β-Actin was used as a loading control. Values are mean (SD) representative of 3 biological replicates. (b) Volcano Plot (t-test) showing gene expression analysis of hnRNP M KD RNA-Seq data from uninfected cells. X-axis shows fold-change of gene expression and Y-axis shows statistical significance. Down-regulated genes are plotted on the left and up-regulated genes are on the right. Dots highlighted in red show genes included in our analysis. (c) Gene expression analysis of hnRNP M -KD cells compared to SCR control for uninfected cells. Each column represents a biological replicate. Genes that are down-regulated are shown in red and up-regulated genes are show in blue. (d) Gene expression analysis of hnRNP M KD cells compared to scramble control for *Salmonella* infected cells. Each column represents a biological replicate. Genes that are down-regulated are shown in red and up-regulated genes are show in blue. (e) Ingenuity Pathway Analysis of gene expression changes in uninfected and *Salmonella* infected cells. (f) Innate immune gene expression analysis highlighting specific innate immune response genes in *Salmonella* infected hnRNP M KD cells compared to scramble control. Each column represents a biological replicate. (g) RT-qPCR of *Rnf26, Rnf128*, and *Slc6a4* in uninfected hnRNP M KD cells. (h) RT-qPCR of mature *IL6, Mx1*, and *Gbp5* transcripts in Salmonella infected hnRNP M KD cells at 2h and 4h postinfection. (i) RT-qPCR of *IL-1β* and *Tnfα* transcripts in Salmonella infected hnRNP M KD cells at 4 hours post-infection. G, h, and i represent 3 biological replicates where values are means ± SEM, n=3. Statistical significance was determined using two-tailed students’ *t* test. *P < 0.05, **P < 0.01, ***P < 0.001.

We next infected hnRNP M KD1 and SCR cell lines with *Salmonella* Typhimurium at an MOI of 10 and performed RNA-seq analysis on total polyA+ selected RNA collected from uninfected and infected cells at 4h post-infection. We chose 4h as a key innate immune time point as one would expect robust transcriptional activation downstream of TLR4 through both MyD88 and TRIF adapters by 4h^31^. To determine how loss of hnRNP M affected gene expression in both uninfected and *Salmonella*-infected RAW 264.7 macrophages, we used CLC Genomics Workbench differential expression pipeline to identify genes whose expression was regulated by hnRNP M. A number of genes were differentially expressed in uninfected hnRNP M KD cells when compared to SCR control cells, with 391 genes up-regulated and 174 down-regulated (Fig. 1b and 1c) and similar ratios of up- and down-regulated genes were seen in hnRNP M KD cells infected with *Salmonella* compared to SCR control (Fig. 1d). In both conditions (+/− *Salmonella*), loss of hnRNP M led to far more upregulated genes (blue) than downregulated (red), indicating that hnRNP M generally acts as a repressor of gene expression, consistent with previous reports of it repressing pre-mRNA splicing^32,33^. Interestingly, we observed only 25% overlap between genes that were differentially expressed in uninfected and *Salmonella*-infected macrophages, suggesting that hnRNP M has distinct modes of operation depending on the activation state of a macrophage (Fig. S1). Unbiased canonical pathways analysis revealed strong enrichment for differentially expressed genes in innate immune signaling pathways in *Salmonella*-infected hnRNP M KD cells (Fig. 1e) and manual analysis of these lists revealed a number of important chemokines (e.g. *Cxcl16, Ccl17, Ccl2, Ccl7*), antiviral molecules (e.g. *Isg15, Mx1)*, and pro-inflammatory cytokines (e.g. *IL6, Mip-1α (Ccl3), IL-18)* whose expression were dramatically affected by loss of hnRNP M (Fig. 1f). Additional pathways enriched for hnRNP M-dependent genes can be found in Fig. S1B and a list of all impacted genes (+/− 1.5 fold change) can be found in Table S1.

To validate the RNA-SEQ gene expression changes, we used RT-qPCR to measure transcript levels of genes from both lists (uninfected SCR vs. hnRNP M KD and *Salmonella*-infected SCR vs. hnRNP M KD). We confirmed overexpression of several genes in uninfected hnRNP M KD cells (*Rnf26, Rnf128, Slc6a4*; Fig. 1g), as well as hyperinduction of genes in hnRNP M KD cells 2h and 4h *post-Salmonella* infection (*IL6, Mx1, Gbp5, Adora2a*, and *Marcks*) (Fig. 1h and S1). Importantly, we found that induction of other pro-inflammatory mediators such as *IL1β* and *Tnfα* did not rely on hnRNP M (Fig. 1i), suggesting that hnRNP M’s ability to regulate gene expression is conferred by specificity at the individual transcript level, rather than being common to a transcriptional regulon (e.g. NFκB, IRF3, STAT1). Together, these results reveal a previously unappreciated role for hnRNP M in repressing specific innate immune transcripts in macrophages.

### hnRNP M regulates a specific subset of innate immune genes upon diverse innate immune stimuli

*Salmonella* encodes several pathogen-associated molecular patterns (PAMPs) that serve as potent activators of pattern recognition receptors. *Salmonella* can also activate pro-inflammatory gene expression via its virulence-associated type III secretion system ^34^. To begin to determine the nature of the signal through which hnRNP M-dependent gene expression changes occur, we first tested whether LPS, a potent agonist of TLR4^35^ and component of the *Salmonella* outer membrane, was sufficient to hyperinduce *IL6* expression (Fig. 2a). SCR control and hnRNP M KD cell lines were treated with 100 ng/mL of LPS, and *IL6* mRNA levels were measured by RT-qPCR. Similar to *Salmonella* infection, we observed a 3-4-fold hyperinduction of *IL6* in hnRNP M KD cells treated with LPS for 2h and 4h, confirming that transcriptional activation of *IL6* occurs downstream TLR4 (Fig. 2b). Importantly, hyperinduction of *IL6* mRNA in both LPS-treated and *Salmonella*-infected hnRNP M KD macrophages increased IL-6 protein levels 3-6 fold (Fig. 2c), indicating that hnRNP M repression of *IL6* mRNA processing impacts protein outputs in a biologically meaningful way. We believe hnRNP M mainly functions to repress *IL6* expression early in macrophage activation, as we did not observe a difference in *IL6* mRNA levels between SCR and hnRNP M KD at later time points post-LPS treatment (Fig. S2).

**Figure 2.**
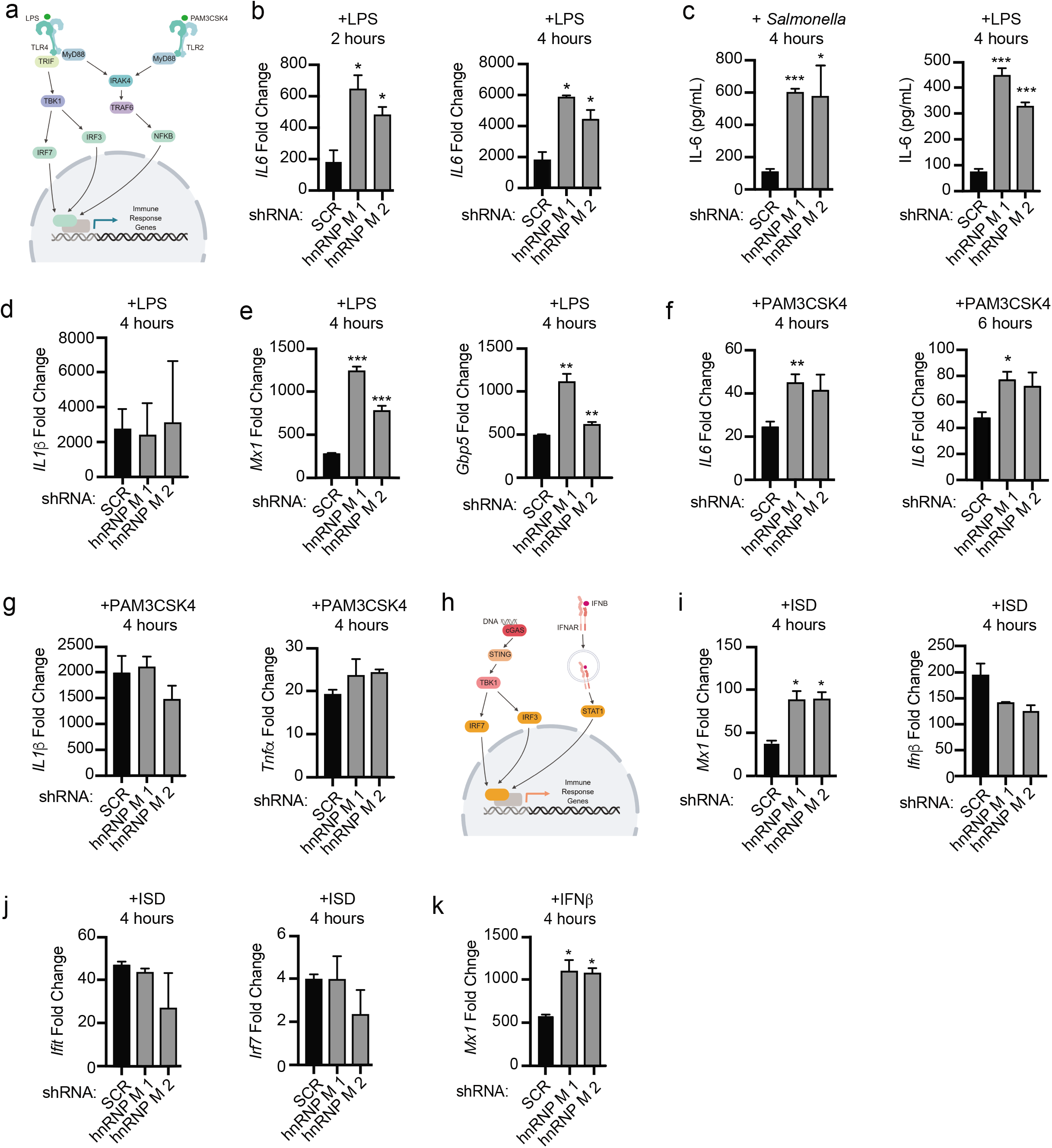
hnRNP M-dependent regulation of innate immune gene expression occurs downstream of multiple innate immune stimuli. (a) Model of TLR4 and TLR2 signaling. (b) RT-qPCR of *IL6* mRNA levels in SCR control and hnRNP M KD cells treated with LPS for 2 hours and 4 hours. (c) IL-6 ELISA with supernatants collected 4 hours post-infection and post-LPS treatment. (d) RT-qPCR of *IL-1β* transcripts in LPS treated hnRNP M KD cells at 4 hours postactivation. (e) RT-qPCR of *Mx1* and *Gbp5* mRNA levels in SCR control and hnRNP M KD cells treated with LPS for 4 hours. (f) RT-qPCR of mature *IL6* levels in SCR control and hnRNP M KD cells treated with PAM3CSK4 for 4 hours and 6 hours. (g) RT-qPCR of *IL-1β* transcripts in PAM3CSK4 treated hnRNP M KD cells at 4 hours post-activation. (h) Model of cGAS-mediated cytosolic DNA sensing and IFNAR signaling. (i) RT-qPCR of *Mx1* and *Ifnβ* mRNA levels in SCR control and hnRNP M KD cells at 4h following ISD transfection. (j) RT-qPCR of interferon-stimulated genes (*Ifit* and *Irf7*) in SCR control and hnRNP M KD cells at 4h following ISD transfection (k) RT-qPCR of *Mx1* transcript in SCR control and hnRNP M KD cells treated with IFN-β for 4 hours. b, c, d, e, f, g, i, j, and k represent 3 biological replicates where values are means ± SEM, n=3. Statistical significance was determined using two-tailed students’ *t*test. *P < 0.05, **P < 0.01, ***P < 0.001.

Consistent with our RNA-SEQ and RT-qPCR data from *Salmonella*-infected cells, *IL1β* (Fig. 2d) and *Tnfα* (Fig. S2), showed no changes in expression after LPS-treatment in hnRNP M KD cells, despite being tremendously upregulated. On the other hand, both *Mx1* and *Gbp5* were hyperinduced in hnRNP M KD cells after LPS treatment (Fig. 2e). Rather than being regulated by NF-κB, transcription of *Mx1* and *Gbp5* is activated by STAT1 downstream of IFNβ signaling, following IFNβ expression via the TRIF/IRF3 axis (Fig. 2h). These results hinted at a mechanism for hnRNP M-dependent repression that is independent of transcription factor specificity and is instead dependent on individual transcripts.

To more directly test the idea that hnRNP M’s target specificity is at the level of the transcript itself, we tested whether these same genes (*IL6* or *Mx1*) were hyperinduced in hnRNP M KD cells treated with a panel of innate immune agonists. Treatment with 100 ng/ml of the TLR2/1 agonist PAM3CSK4 hyperinduced *IL6* expression in hnRNP M KD cells compared to SCR controls (Fig. 2f), while *Tnfa* and *IL1β* mRNA levels remained similar (Fig. 2g). Likewise, transfection of hnRNP M KD cells with 1 μg/mL interferon stimulatory DNA (ISD), a potent agonist of cytosolic DNA sensing and IRF3-mediated transcription downstream of the cGAS/STING/TBK1 axis ^36^(Fig. 2h) led to hyperinduction of *Mx1* in hnRNP M KD cells (Fig. 2i), whereas *Ifnβ* and other interferon-stimulated genes (ISGs) regulated by IRF3 (*Ifit1* and *Irf7*) were expressed at similar levels (Fig. 2i and 2j). Direct engagement of the interferon receptor (IFNAR) with recombinant IFNβ also resulted in *Mx1* hyperinduction in hnRNP M KD cells (Fig. 2k). Collectively, these results bolster a model whereby hnRNP M represses mRNA expression of a specific subset of innate immune genes, regardless of how those genes are induced.

### hnRNP M influences gene expression outcomes at the level of pre-mRNA splicing

Previous studies of hnRNP M’s role in gene expression have shown that it can enhance or silence splicing of alternatively spliced exons^32,37–39^. However, because hnRNPs are known to regulate post-transcriptional gene expression at many steps (e.g. mRNA decay, mRNA export, translational repression), we first asked whether loss of hnRNP M could specifically influence constitutive intron removal and/or alternative splicing in LPS-activated macrophages. We chose *IL6* as a model transcript because: (1) it has a simple intron-exon architecture for a mammalian gene, with four relatively short introns (165, 1271, 3059 and 1226 nucleotides, respectively); (2) it was significantly and robustly hyperinduced by loss of hnRNP M (Fig. 1f and 1h); and (3) it is a crucial component of the macrophage inflammatory response. Using RT-qPCR, we first measured the relative abundance of each *IL6* intron-exon junction (Fig. 3a) in SCR control cells to assess how intron removal proceeded on *IL6* pre-mRNAs in cells containing hnRNP M. Primers were designed so as to only amplify introns that are still part of pre-mRNAs and not released intron lariats. At two hours post-LPS treatment, most of the *IL6* transcripts we detected are partially processed, with intron 1 and to some extent intron 4 being preferentially removed and introns 2 and 3 being retained (Fig. 3b). We then compared the relative abundance of *IL6* introns in SCR control cells to those in hnRNP M KD macrophages and observed a dramatic and specific decrease in intron 3-containing *IL6* pre-mRNAs in the absence of hnRNP M. This decrease in *IL6* intron 3 starkly contrasted other *IL6* intron-exon and exon-exon junctions, which were overall more abundant in the absence of hnRNP M (Fig. 3c). The fact that we observe vastly different amounts of intron 2 and 4-containing *IL6* pre-mRNAs compared to intron 3-containing *IL6* pre-mRNAs in hnRNP M KD macrophages speaks against hnRNP M impacting *IL6* expression transcriptionally and instead argues strongly for the protein playing a role in *IL6* pre-mRNA processing. These data demonstrate that in *IL6* pre-mRNAs accumulate in the absence of hnRNP M and suggest that *IL6* intron 3 plays a privileged role in dictating the maturation of *IL6* mRNAs. Based on this result, we propose that hnRNP M is important for controlling constitutive splicing of certain introns in macrophages and that it can inhibit removal of intron 3 from *IL6* pre-mRNA in order to limit accumulation of the fully spliced mRNA.

**Figure 3.**
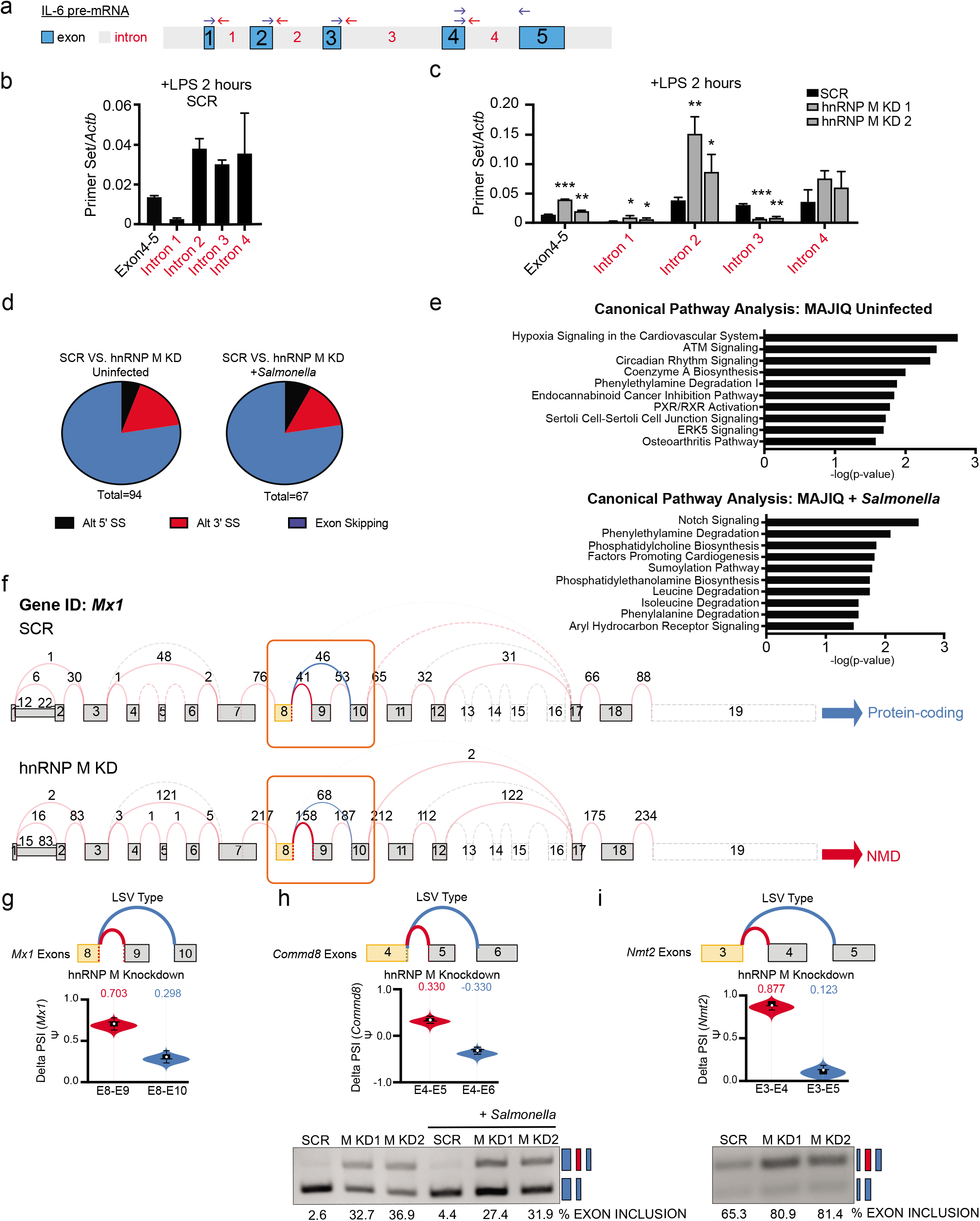
hnRNP M influences gene expression outcomes at the level of pre-mRNA splicing (a) Diagram of *IL6* pre-mRNA with introns (gray) and exons (blue). (b) RT-qPCR of *IL6* exon-exon and intron-exon junctions in SCR control macrophages at 2h post-LPS treatment. (c) RT-qPCR of *IL6* exon-exon and intron-exon junctions in SCR vs. hnRNP M KD1 and KD2 at 2h post-LPS treatment. (d) Categorization of alternative splicing events identified via MAJIQ in uninfected SCR vs. hnRNP M KD1 samples and in *Salmonella*-infected SCR vs. hnRNP M KD1 samples. (e) Ingenuity Pathway Analysis of genes from MAJIQ analysis in uninfected and *Salmonella*-infected cells. (f) VOILA output of *Mx1* transcript model in SCR and hnRNP M KD1 cells infected with *Salmonella.* (g) Violin plots depicting alternative splicing of *Mx1.* (h) Violin plots depicting alternative splicing of *Commd8* with quantification of RT-PCR results. (i) Violin plots depicting alternative splicing of *Nmt2* with quantification of RT-PCR results. B and c are representative of two independent experiments that showed the same result with values representing means (SD), n=3. Statistical significance was determined using two-tailed students’ *t* test. *P < 0.05, **P < 0.01, ***P < 0.001, ****P < 0.0001, D was done with MAJIQ gene outputs from RNA-seq samples containing 3 biological replicates. H and i are representative of 3 biological replicates. For exon inclusion values, n=1.

We next wanted to explore if loss of hnRNP M also influenced alternative splicing in uninfected and *Salmonella*-infected macrophages. To do so, we employed an algorithm for local splice variation (LSV) analysis called MAJIQ (modeling alternative junction inclusion quantification)^40^. MAJIQ allows identification, quantification, and visualization of diverse LSVs, including alternative 5’ or 3’ splice site usage and exon skipping, across different experimental conditions. MAJIQ identified a total of 94 LSVs in uninfected SCR vs. hnRNP M KD macrophages and 67 LSVs in *Salmonella*-infected SCR vs. hnRNP M KD macrophages (probability [| delta PSI|, ≥20%], >95%) (Fig. 3d). The vast majority of the LSVs identified in SCR vs. hnRNP M KD cells were exon skipping events. Subsequent visualization of these LSVs by VOILA analysis revealed that loss of hnRNP M generally correlated with increased exon inclusion in both uninfected and *Salmonella*-infected macrophages. In other words, the presence of hnRNP M led to more exon skipping, which is consistent with a role for hnRNP M in splicing repression. We also conducted IPA pathway analysis of alternatively spliced transcripts to identify pathways enriched for hnRNP M-dependent changes. In contrast to our global gene expression IPA analysis, we observed no enrichment for genes in innate immune-related pathways in either uninfected or *Salmonella*-infected macrophages (Fig. 3e). In fact, only 3 transcripts had both significant expression changes (via RNA-SEQ) and significant delta PSI changes (via MAJIQ), suggesting that hnRNP M’s role in influencing steady state gene expression of innate immune transcripts is distinct from its role in controlling alternative splicing decisions (Fig. S3).

*Mx1*, an anti-viral GTPase, was one of the three transcripts significantly impacted by loss of hnRNP M at the levels of gene expression (Fig. 1f & 1h) and alternative splicing (Fig. 3f). Specifically, MAJIQ identified an exon inclusion event of *Mx1* “exon 9” that was significantly more frequent in hnRNP M KD uninfected macrophages vs SCR control uninfected macrophages (delta PSI exon 8-exon 9 = 0.703 vs. exon 8-exon 10 = 0.298) (Fig. 3g). Inclusion of this exon 9 introduces a premature stop codon and exon 9-containing transcript isoforms of *Mx1* are annotated as nonsense mediated decay targets. Therefore, the overall abundance of MX1 protein may be regulated by hnRNP M at multiple post-transcriptional processing steps, i.e. bulk transcript abundance and proportion of functional protein-encoding transcripts. MAJIQ also reported increased exon inclusion events for *Commd8*, a putative transcriptional regulator, and *Nmt2*, an N-myristoyltransferase, and we confirmed each of these LSVs by semiquantitative RT-PCR (Fig. 3h & Fig. 3i). Collectively, these data illustrate that hnRNP M can repress splicing of both constitutive and alternative introns, leading to distinct gene expression outcomes and protein synthesis outcomes in macrophages.

### hnRNP M is enriched at the level of chromatin and at the IL6 genomic locus

To get a better understanding of how hnRNP M controls pre-mRNA splicing, we next asked where hnRNP M localized in RAW 264.7 macrophages and whether its localization changed upon TLR4 activation. Other hnRNP family members have been found to translocate to the cytoplasm in response to several different types of stimuli including Vesicular stomatitis virus (VSV) infection, osmotic shock, and inhibition of transcription^14,41,42^. In particular, previous reports of hnRNP U have shown that it shuttles out of the nucleus following LPS treatment of macrophages^10^, and hnRNP M itself has been shown to translocate from the nucleus to the cytoplasm during enterovirus infection of HeLa cells^10,28^. Based on our data implicating hnRNP M in splicing, we predicted that it can function in the nucleus and indeed, several algorithms including NLS Mapper^43^ and PredictProtein^44^ predicted hnRNP M is a predominantly nuclear protein (NLS Mapper score 8.5/10; PredictProtein 98/100) (Fig. 4a).

**Figure 4.**
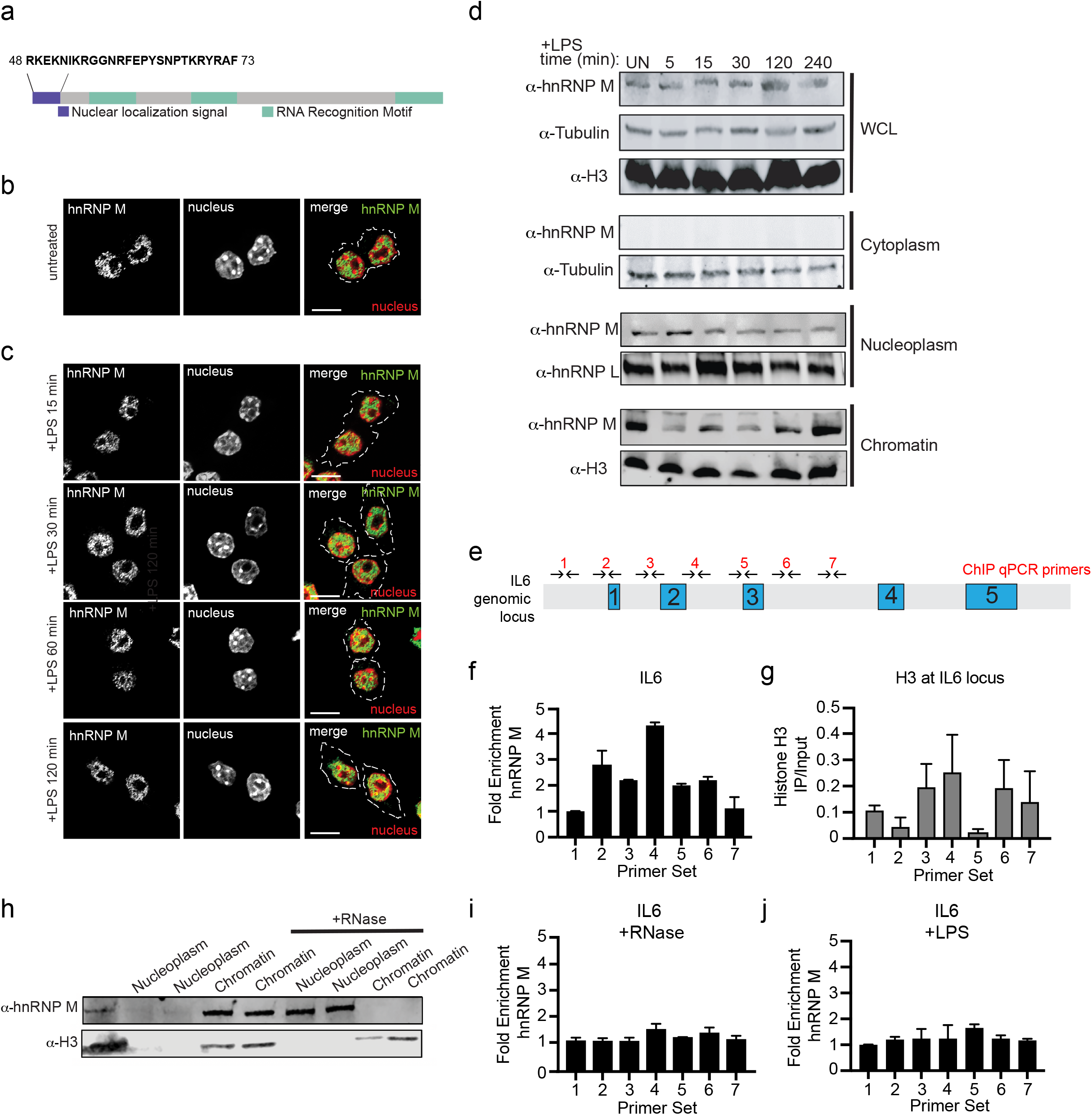
hnRNP M is a nuclear protein that associates with the IL-6 genomic locus in an RNA-dependent fashion (a) Schematic diagram of hnRNP M, highlighting the nuclear localization signal (purple) and three RNA Recognition Motifs (green). (b) Immunofluorescence images of uninfected RAW 264.7 macrophages immunostained with anti-hnRNP M (green). (c) Immunofluorescence images of RAW 264.7 macrophages stimulated with LPS for the respective time points and immunostained with anti-hnRNP M (green). (d) Western blot analysis of cellular fractions with anti-hnRNP M and loading controls of cytoplasm (tubulin), nucleoplasm (hnRNP L) and chromatin (H3) fractions of uninfected and LPS stimulated RAW 264.7 macrophages. (e) CHIP-qPCR primers designed to tile *IL6* locus. (f) RT-qPCR of ChIP at the *IL6* genomic locus with anti-hnRNP M in resting RAW 264.7 macrophages. (g) RT-qPCR of histone H3 CHIP at the *IL6* genomic locus in RAW 264.7 macrophages. (h) Western blot analysis of nuclear and chromatin fractions with anti-hnRNP M and Histone H3 (control) with untreated and RNase treated nuclear fractions. (i) ChIP-qPCR with anti-hnRNP M in RAW 264.7 macrophages treated with RNase. (j) ChIP qPCR of hnRNP M in macrophages treated with 100ng/mL LPS for 1h. F and g values are means ± SEM representative of 2 biological replicates, n=2. I and j values are means ± SEM representative of 3 biological replicates, n=3. Statistical significance was determined using two-tailed students’ *t* test. *P < 0.05, **P < 0.01, ***P < 0.001.

To examine hnRNP M localization, we performed immunofluorescence microscopy in uninfected macrophages using an antibody that detects endogenous hnRNP M and observed significant enrichment of hnRNP M in the nucleus (Fig. 4b). We next treated macrophages with LPS and analyzed hnRNP M localization at various timepoints to determine how activation might alter localization, but we observed no major changes to hnRNP M localization. (Fig. 4c). This was true for both endogenous hnRNP M and a 3xFLAG-hnRNP M allele stably expressed in macrophages (Fig. S4). As a control, we monitored the translocation of hnRNP U upon LPS treatment and observed nuclear to cytoplasmic translocation, consistent with previous reports (Fig. S4). Based on these results, we concluded that hnRNP M is a nuclear protein in macrophages and that LPS treatment does not trigger translocation to another cellular compartment.

We next sought to understand more precisely where in the nucleus hnRNP M was enriched since intron recognition and removal can occur at the level of chromatin, while nascent transcripts are still tethered to RNA Polymerase II^1,45–47^, or in the nucleoplasm after pre-mRNAs have been fully transcribed and released. To this end, we performed a cellular fractionation experiment in RAW 264.7 macrophages over a time course of LPS treatment and visualized hnRNP M localization via western blot (Fig. 4d). Consistent with our immunofluorescence experiments, we did not detect hnRNP M in the cytoplasmic fraction at any time point. However, we observed hnRNP M in both the nucleoplasm and the chromatin over the LPS treatment. Macrophages stably expressing 3xFLAG-hnRNP M showed a similar hnRNP M distribution between the nucleoplasm and chromatin (Fig. S4). We did not observe significant redistribution of either endogenous or 3xFLAG-hnRNP M between the nucleoplasm and chromatin fractions upon LPS treatment (Fig. 4c and S4). Together, fractionation and immunofluorescence experiments confirm that a population of hnRNP M associates with chromatin and the protein does not grossly redistribute in the cell upon LPS treatment.

### hnRNP M’s association with the IL6 locus is RNA dependent and controlled by TLR4 signaling

We next wanted to determine if hnRNP M’s association with chromatin was specific for the genomic loci of genes whose regulation was impacted by hnRNP M (Fig. 1f). We hypothesized that if hnRNP M repression of *IL6* intron 3 removal occurs at the nascent transcript level, hnRNP M may associate with the *IL6* genomic locus. To test this, we performed chromatin immunoprecipitation (ChIP)-qPCR. ChIP has been used extensively in yeast, and to some extent in mammals, as a spatiotemporal read out of splicing factor recruitment to nascent transcripts^48–52^. To test whether hnRNP M associated with the *IL6* genomic locus, we immunoprecipitated endogenous hnRNP M from untreated macrophages and determined its association with the *IL6* locus (DNA), using a series of tiling primers spaced approximately 500 bp apart (Fig. 4e). We observed no enrichment of hnRNP M in the promoter region of *IL6*, consistent with it playing a mainly post-transcriptional role in *IL6* processing (Fig. 4f, primer set 1). We did however, observe significant enrichment of hnRNP M at several primer sets in the *IL6* gene, most notably over the intron 2-intron 3 region (Fig. 4f, primer set 4). Previously published CLIP-seq experiments identified a GUGGUGG consensus site for hnRNP M; such a site exists in intron 2 of *IL6* and several similar motifs are found in *IL6* intron 3 (Fig. S4). ChIP-qPCR of histone H3, which showed clear depletion of nucleosomes around the *IL6* transcription start site (primer sets 1 & 2), was performed to control for genomic DNA accessibility and/or primer set efficiency (Fig. 4g). Together, these results reveal that hnRNP M can associate with the genomic locus of genes like *IL6* whose splicing it represses, suggesting that it functions co-transcriptionally, prior to release of transcripts from RNA polymerase II.

Because hnRNP M is an RNA binding protein by definition, the next question we asked was whether its association with the *IL6* genomic locus was dependent on RNA or occurred through interactions with chromatin-associated proteins. To this end, we performed another cellular fractionation experiment with and without RNase A digestion^52^. We observed a dramatic redistribution of hnRNP M from the chromatin to the nucleoplasm following addition of RNase, providing strong evidence that its association with chromatin is through RNA (Fig. 4h). These data are consistent with a previously published screen of splicing factors where hnRNP M was found to have cell type-specific RNA-dependent chromatin association^53^. Based on these results looking at bulk chromatin, we next set out to test whether hnRNP M’s association with the *IL6* genomic locus was similarly RNA-dependent. We performed ChIP-qPCR as above but with an additional RNase treatment after sonication for 30 min at 37°C. RNase treatment completely abolished any enrichment of hnRNP M at the *IL6* locus, confirming that its association with the *IL6* gene depends on RNA (Fig. 4i).

If hnRNP M acts as a repressor of *IL6* splicing by binding to nascent transcripts at the *IL6* locus, we hypothesized that this repression might be relieved upon TLR4 activation, which would allow a cell to robustly induce *IL6* expression following pathogen sensing. To test this, we performed ChIP-qPCR of hnRNP M at the *IL6* locus in RAW 264.7 macrophages treated with LPS for 1h. Remarkably, we observed a complete loss of hnRNP M enrichment at all primer sets along the *IL6* gene body, including those over intron 2 and 3, following LPS treatment (Fig. 4j). This result strongly links hnRNP M’s ability to repress *IL6* with its presence at the *IL6* genomic locus and suggests that TLR4 signaling controls hnRNP M’s repressor activity.

### Phosphorylation of hnRNP M at S574 downstream of TLR4 activation controls its ability to repress expression of innate immune transcripts

A recently published phosphoproteomics dataset identified a number of splicing factors that were differentially phosphorylated during infection with the intracellular bacterium *Mycobacterium tuberculosis*^20^. Because it is not a gram-negative bacterium, *M. tuberculosis* does not activate TLR4 via LPS, but it does express surface peptidoglycan, which is an agonist of TLR2. Having confirmed hnRNP M-dependent regulation of *IL6* following treatment with a TLR2 agonist (PAM3CSK4) (Fig. 2f), we reasoned that TLR2 activation upon *M. tuberculosis* infection may lead to the same changes in hnRNP M phosphorylation as would TLR4 activation during *Salmonella* infection. We thus leveraged the *M. tuberculosis* global phosphoproteomics dataset from Penn *et al.*^20^, identified 5 differentially phosphorylated serine residues on hnRNP M (S85, S431, S480, S574, and S636) (Fig. 5a), and generated 3xFLAG-hnRNP M constructs with phosphomimic (S→D) or phosphodead (S→A) mutations at each of the serines. We then made stable RAW 264.7 macrophages expressing each of these alleles (Fig. 6b) in wild-type RAW 264.7 macrophages containing a wild-type hnRNP M allele and measured *IL6* and *Mx1* expression 4 hours *post-Salmonella* infection. While overexpression of 3xFLAG-hnRNP M itself did not have a major impact on *IL6* or *Mx1* induction, expression of hnRNP M S574D, a phosphomimetic allele, led to hyperinduction of these transcripts, essentially phenocopying hnRNP M KD cells (Fig. 5b & 5c). Several other phosphomutant alleles (S85D, red bar, S431A, light blue bar, S480A/D, green bars) also affected *IL6* and *Mx1* induction but to a much lesser extent (Fig. 5b and 5c). Interestingly, mutating S587, which is a repeat of the S574-containing sequence (MGANS(ph)LER), did not affect the regulation of *IL6* or *Mx1* (Fig. S5), suggesting the location of these serines is critical and that phosphorylation-dependent regulation of hnRNP M is specific for select serine residues (Fig. S5).

**Figure 5.**
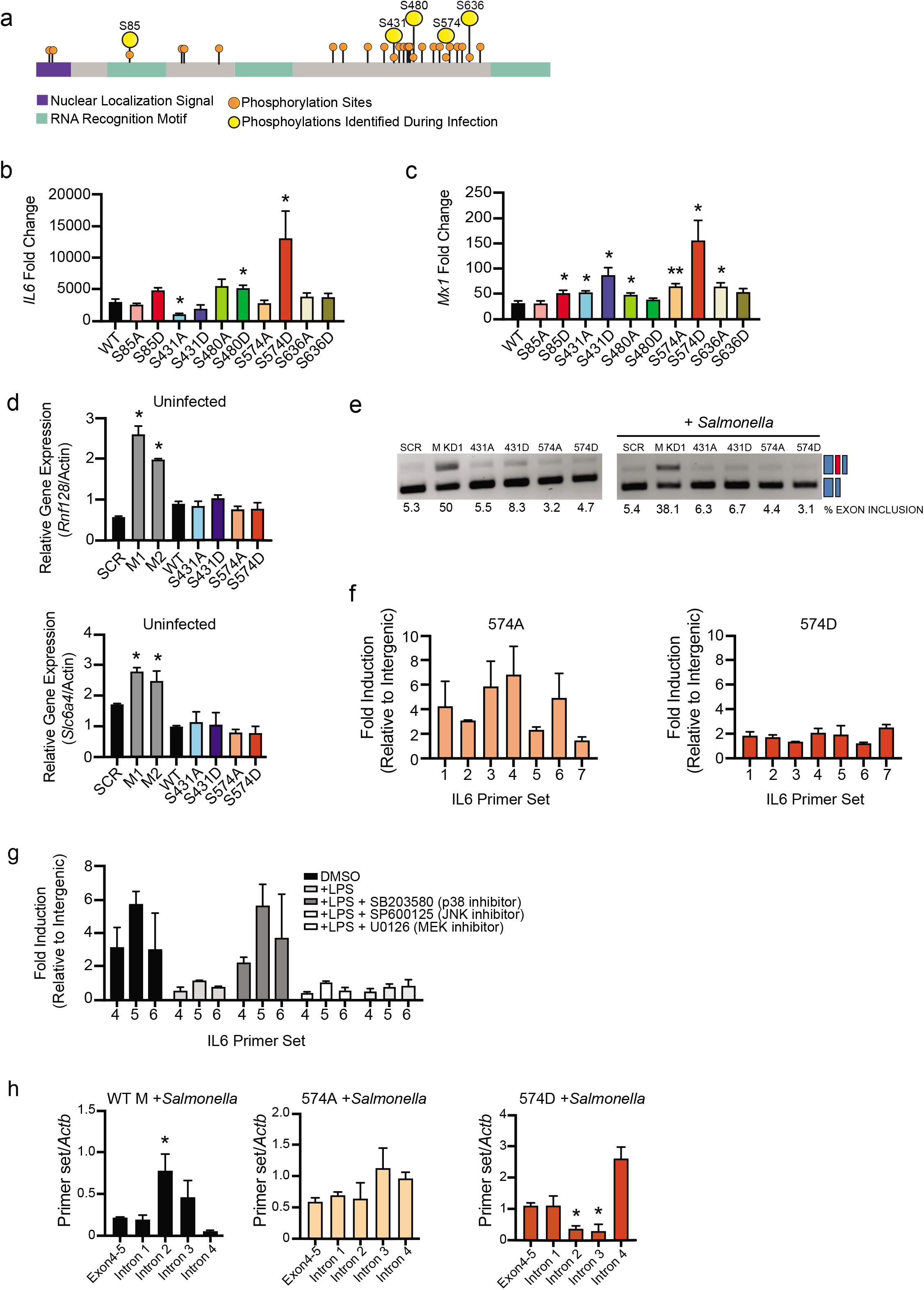
Phosphorylation of hnRNP M at S574 downstream of TLR4 activation controls its ability to repress expression of innate immune transcripts (a) Protein diagram of hnRNP M indicating location of phosphorylation sites identified by SILAC/mass spectrometry with nuclear localization signal shown in purple and RNA-Recognizing Motif (RRM) shown in in green. (b) RT-qPCR of mature *IL6* in WT hnRNP M-FLAG and phosphomutants in macrophages infected with *Salmonella* 4 hours post-infection. (c) RT-qPCR of *Mx1* in WT hnRNP M-FLAG and phosphomutants in macrophages infected with *Salmonella* 4 hours post-infection. (d) RT-qPCR of *Rnf128* and *Slc6a4* in uninfected, WT 3xFL-hnRNP M, and phosphomutants. (e) Semiquantitative PCR of *Commd8* alternative splicing in cells expressing SCR or hnRNP M KD constructs alongside phosphomutant-expressing alleles. (f) ChIP-qPCR of hnRNP M-S574A/D alleles at the *IL6* genomic locus. (g) ChIP-qPCR of wild-type 3xFL-hnRNP M in the presence of 100ng/mL LPS and various MAPK inhibitors (SB203580, SP600125, and U0126). (h) RT-qPCR of *IL6* intron-exon junctions and the exon 4-5 mature junction in 3xFL-hnRNP M, 3xFL-hnRNP M 574A and 574D phosphomutants, in macrophages infected with *Salmonella* 4 hours postinfection. B, c, and d are representative of 3 biological replicates with values indicating means ± SEM, n=3. F g, and h are representative of 2 biological replicates values indicating means ± SEM, n=2. I is representative of 2 independent experiments that showed the same result with values representing means (SD), n=3. Statistical significance was determined using two-tailed students’ *t* test. *P < 0.05, **P < 0.01, ***P < 0.001.

**Figure 6.**
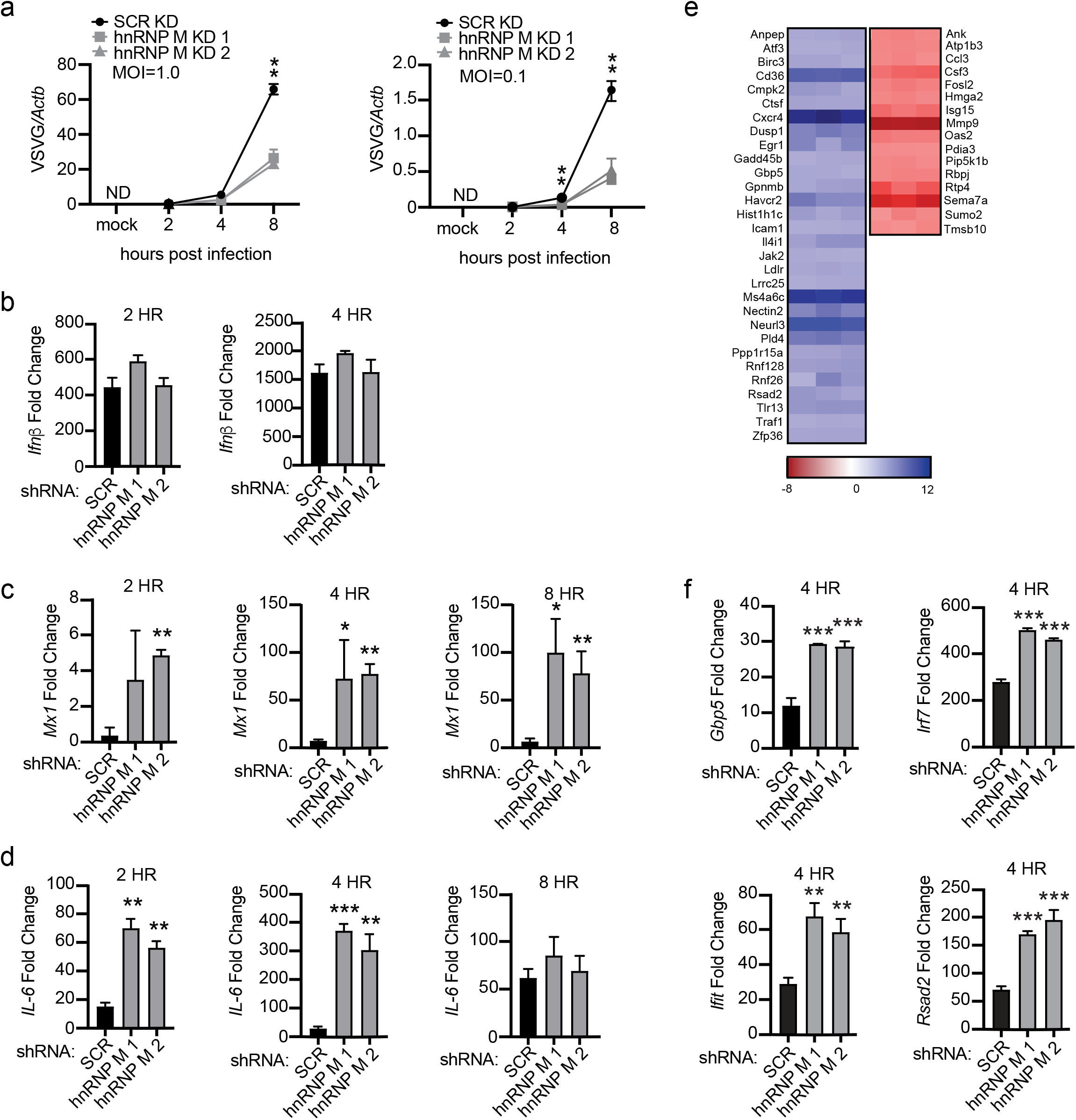
Knockdown of hnRNP M enhances macrophage’s ability to control viral infection. (a) Viral replication in hnRNP M KD and SCR control RAW 264.7 macrophages infected with VSV (MOI=1.0, MOI=0.1, or Mock) at 2h, 4h, and 8h post-infection. (b) RT-qPCR of *Ifnβ* mRNA levels in SCR control and hnRNP M KD cells at 2h and 4h post-infection, MOI=1. (c) RT-qPCR of *Mx1* transcript in VSV-infected SCR control and hnRNP M KD cells at 2h, 4h, and 8h postinfection, MOI=1. (d) RT-qPCR of *IL6* transcript in SCR control and hnRNP M KD cells at 2h, 4h, and 8h post-infection MOI=1. (e) Differential gene expression in hnRNP M KD cells compared to SCR control macrophages from earlier RNA-SEQ analysis (Fig. 1) highlighting known viral response genes. (f) RT-qPCR of *Ifit, Irf7, Rsad2*, and *Gbp5* mRNA levels in SCR control and hnRNP M KD cells at 4h post-infection, MOI=1. A, b, c, d, and e are representative of 2 biological replicates with values indicating means ± SEM, n=2. F is representative of 2 independent experiments that showed the same result with values representing means (SD), n=3. Statistical significance was determined using two-tailed students’ *t*test. *P < 0.05, **P < 0.01.

Having implicated hnRNP M phosphorylation at S574 in controlling *IL6* and *Mx1* expression, we next wanted to see how phosphorylation affected transcripts whose expression in uninfected cells was higher in the absence of hnRNP M (Fig. 1g). While we again observed elevated expression of these transcripts in the absence of hnRNP M (hnRNP M KD, grey bars), introduction of the phosphomutant alleles (S431A/D and S574A/D) had no effect on *Rnf128, Rnf26*, or *Slc6a4* transcript levels (Fig. 5d). Expression of these genes was similarly unaffected by the other hnRNP M phosphomutants (Fig. S5). Alternative splicing of *Commd8* was also unaffected by any of the phophosmutants in either uninfected or *Salmonella*-infected cells (Fig. 5e). Together, these data provide strong evidence that hnRNP M’s ability to regulate the expression of constitutively expressed genes and/or influence alternative splicing decisions does not rely on phosphorylation at 574, whereas its role in regulating innate immune transcripts induced during infection is specifically controlled by this post-translational modification downstream of pathogen sensing.

To further examine how these phosphorylated residues contributed to hnRNP M’s function at an LPS-induced gene like *IL6*, we first performed cellular fractionation to determine the localization of mutant alleles. We found that like wild-type hnRNP M, each hnRNP M phosphomutant was, to some extent, enriched in the chromatin in untreated cells (Fig. S5). However, in ChIP experiments looking specifically at the *IL6* locus, the S574D phosphomimic allele displayed virtually no enrichment compared to 574A phosphodead allele, whose enrichment profile was similar to that of wild-type hnRNP M (Fig. 5f and Fig. 4f). Indeed, hnRNP M 574D ChIPs more closely resembled those from RNase or LPS-treated samples (Fig. 4i and 4j). These data point to phosphorylation of residue S574 in controlling hnRNP M’s ability to co-transcriptionally repress processing of chromatin-associated *IL6* pre-mRNAs.

We next sought to better understand how hnRNP M is phosphorylated at these key residues. TLR4 activation sets off a number of signaling cascades, including p38, MEK1/2 (ERK), and JNK MAP kinases. Previous reports have implicated each of these pathways in regulating *IL6* expression downstream of innate immune stimuli^54^, but it is not known if these cascades control splicing factor phosphorylation. To test the role of each cascade in hnRNP M-dependent repression of *IL6*, we performed ChIP experiments in the presence of LPS and specific inhibitors of p38 (SB203580), JNK (SP600125), or MEK (U0126). We again observed LPS-dependent loss of hnRNP M enrichment at *IL6* (primer sets 4, 5, and 6), and treatment with JNK and MEK inhibitors had no effect on hnRNP M release. However, in the presence of the p38 inhibitor, hnRNP M remained associated with the *IL6* genomic locus after LPS treatment (Fig. 5h), demonstrating that p38 signaling promotes release of hnRNP M from the *IL6* genomic locus.

Lastly, to interrogate the mechanism driving *IL6* hyperinduction in hnRNP M 574D-expressing cells, we asked whether *IL6* intron removal was affected by expression of the phosphomutant alleles. Using the same RT-qPCR approach used in Figure 2B, we detected an increase in *IL6* pre-mRNAs containing introns 2 and 3 in macrophages overexpressing a wildtype hnRNP M allele, consistent with hnRNP M slowing *IL6* intron removal. Conversely, these same introns were removed more efficiently in the presence of hnRNP M 574D while no difference was observed in 574A-expressing cells (Fig. 5h). These data strongly support a model whereby phosphorylation of hnRNP M at S574 relieves its ability to act as a splicing repressor, allowing for rapid removal of *IL6* introns and uprgulation of *IL6* mRNA. Together, they demonstrate a novel role for constitutive intron removal in mediating *IL6* expression in macrophages.

### Loss of hnRNP M enhances macrophage’s ability to control viral infection

Because loss of hnRNP M resulted in hyperinduction of a variety of cell-intrinsic antimicrobial molecules and interferon-stimulated genes, we hypothesized that hnRNP M KD cells would be better at controlling viral replication at early time points. To test how loss of hnRNP M influenced viral replication, we infected SCR and hnRNP M KD RAW 264.7 macrophages with VSV, an enveloped RNA virus that can replicate and elicit robust gene expression changes in RAW 264.7 macrophages^55^. Viral replication (levels of VSV-G) was measured over an 8h time course by RT-qPCR in cells infected with a viral MOI of 1 and 0.1. At both MOIs, loss of hnRNP M correlated with dramatic restriction of VSV replication, particularly at the 8h time point (Fig. 6a). As expected, infection with VSV, a potent activator of cytosolic RNA sensing via RIG-I/MAVS^55^, led to robust induction of *Ifnβ* levels in an hnRNP M-independent fashion, as was previously observed in hnRNP M KD cells transfected with cytosolic dsDNA (Fig. 6b and 2i, respectively). Consistent with hnRNP M-dependent regulation occurring downstream of diverse immune stimuli (Fig. 2), VSV-infected hnRNP M KD cells hyperinduced both *IL6* and *Mx1* (Fig. 6c and d).

While *Mx1* itself is a well-characterized anti-viral GTPase, several reports have noted that cell lines like RAW 264.7 that are derived from inbred mouse strains carry mutant, or in some cases non-functional, *Mx1* alleles^56^. Therefore, to begin to predict what hnRNP M-regulated genes may be responsible for enhanced VSV restriction, we manually examined hnRNP M-regulated transcripts in our RNA-SEQ data from resting and *Salmonella*-infected (i.e. TLR4-activated) macrophages and identified a number of genes known to be important for controlling RNA viral replication (Fig. 6e). RT-qPCR confirmed hyperinduction of several antiviral ISGs in hnRNP M KD macrophages at 4h post-VSV infection including *Rsad2* (Viperin), *Ifit, Irf7*, and *Gbp5.* Interestingly, neither *Ifit* nor *Irf7* was identified as an hnRNP M-dependent transcript during *Salmonella* infection, even though both can be expressed downstream of TLR4 through IRF3/IFNAR/STAT1 signaling. This difference may simply reflect kinetic differences in transcript induction following cytosolic RNA sensing vs. TLR4 activation or may indicate that hnRNP M regulates an even broader set of transcripts in macrophages following RNA virus infection. We propose that inhibition of VSV replication in hnRNP M KD macrophages ultimately results from a combination of pro-viral gene downregulation (red genes, Fig. 6e) and anti-viral gene upregulation (blue genes, Fig. 6b and Fig. 6e). Collectively, these data are consistent with hnRNP M playing a critical role in slowing innate immune gene expression and suggest that the presence of hnRNP M can actually blunt macrophage antiviral defenses at early time points following infection with VSV.

## Discussion

Despite the substantial impact pre-mRNA splicing has on gene expression outcomes, little is known about how components of the spliceosome are modified and regulated during cellular reprogramming events, such as macrophage pathogen sensing. Here, we demonstrate that the splicing factor hnRNP M is a critical repressor of a unique regulon of innate immune transcripts (see model in Fig. 7). These transcripts were hyperinduced in hnRNP M KD macrophages downstream of a variety of innate immune stimuli (i.e. *Salmonella* infection, TLR4/TLR2 agonists, recombinant IFN-β, cytosolic dsDNA, RNA virus infection (VSV)) (Fig. 2 and Fig. 6) and hyperinduction of this regulon correlated with enhanced capacity of hnRNP M KD macrophages to control VSV replication at early time points (Fig. 6). We propose that in innate immune cells like macrophages repression of pre-mRNA splicing by hnRNP M serves as a safeguard, dampening the initial ramping up of innate immune gene expression and preventing spurious expression of potent pro-inflammatory molecules in situations where the cell has not fully engaged with a pathogen. The requirement for cells to tightly control expression of potent inflammatory mediators like IL-6 is evidenced by the fact that multiple transcriptional and post-transcriptional mechanisms exist to regulate *IL6*, including chromatin remodeling^57^, mRNA stability^58^, subcellular localization^59^, and now, based on these data, pre-mRNA splicing.

**Figure 7.**
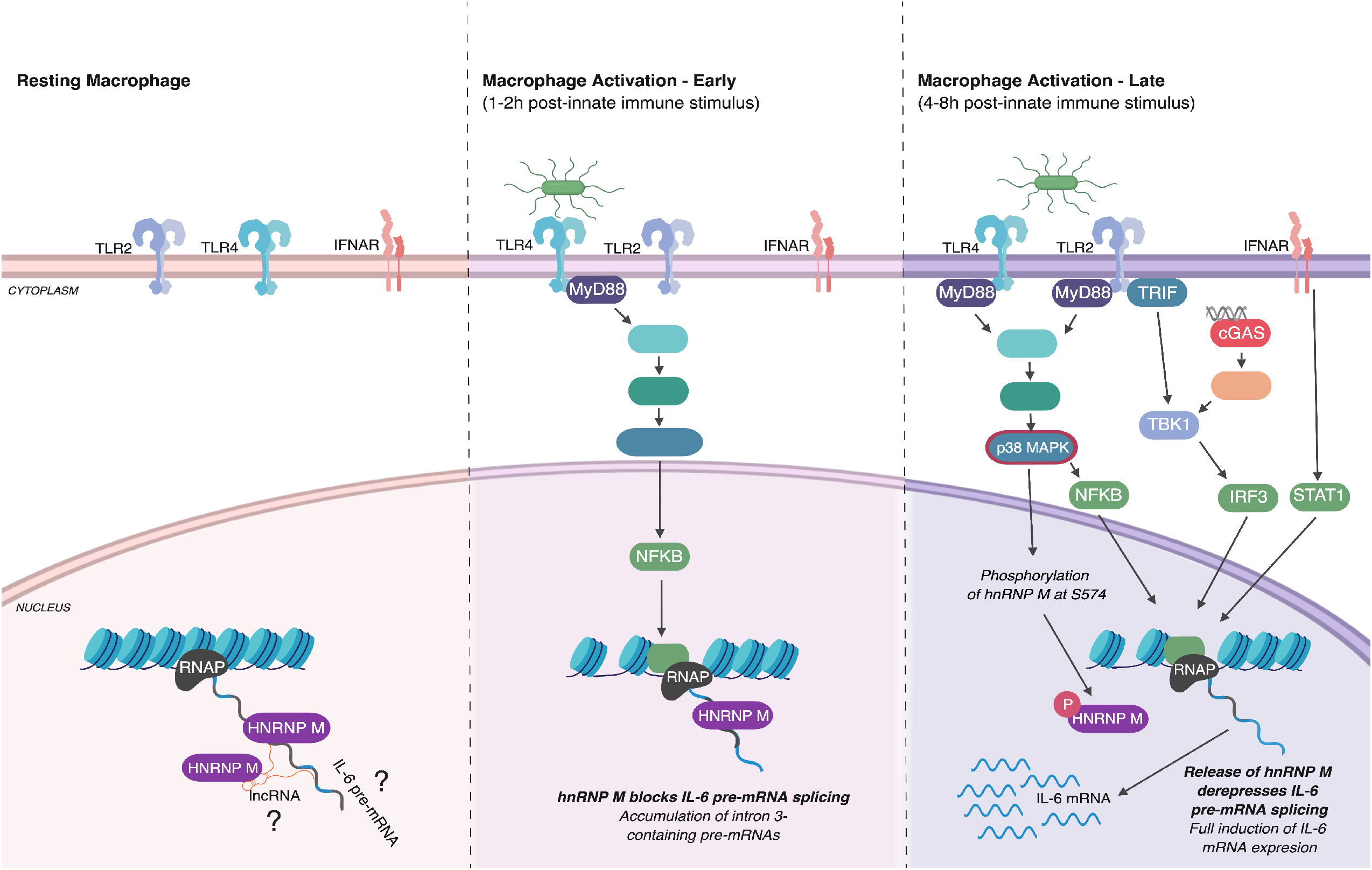
Proposed model for hnRNP M-dependent repression for *IL6* expression in resting, early-, and late-activated macrophages (Left panel) In resting macrophages, hnRNP M associates with chromatin, at the *IL6* genomic locus, through interactions with RNA. These interactions may be direct with target transcripts, or indirect via protein-interactions with other RNA binding proteins or through interactions with other chromatin-associated RNAs, e.g. linc-RNAs. (Middle panel) When macrophages receive an innate immune stimulus, they transcriptionally activate genes like *IL6.* hnRNP M associates with chromatin-bound, pre-mRNAs in these cells, inhibiting *IL6* intron removal and preventing full maturation of *IL6* nascent transcripts. (Right panel) Following extended innate immune activation, hnRNP M is phosphorylated at S574 in a p38-MAPK-dependent fashion. Phosphorylation of hnRNP M releases it from the *IL6* genomic locus, relieves inhibition of *IL6* splicing, and allows for full induction of *IL6* gene expression. Figure generated using Biorender.

Our results point to target transcripts themselves dictating hnRNP M’s specificity, rather than a particular signaling cascade or transcription factor. Data from RNA-SEQ experiments revealed that hnRNP M can activate/repress a cohort of non-inducible “housekeeping” genes in uninfected macrophages and an entirely different set of genes, mainly comprised of innate immune transcripts, following innate immune stimuli. This “switch” in hnRNP M activity is regulated, at least in part, by p38 MAPK-dependent phosphorylation at hnRNP M S574, which impacts the protein’s ability to associate with the *IL6* genomic locus and to repress *IL6* intron removal (Fig. 4, 5, and 7). Our data argue that while hnRNP M influences alternative splicing decisions in resting macrophages, phosphorylation of hnRNP M specifically impacts its ability to regulate a cohort of innate immune transcripts. Based on these observations, we propose that hnRNP M and likely other splicing factors possess distinct capacities for interacting with RNAs and/or proteins depending on how they are post-translationally modified. In this way, innate immune sensing cascades may remodel splicing complexes, for example, by promoting release of hnRNP M from chromatin-associated RNAs via p38-MAPK cascades.

While we do not fully understand the mechanisms driving hnRNP M’s target specificity, our RNA-SEQ data as well as other datasets^1^ demonstrate the presence of cryptic exons in a number of hnRNP M-regulated transcripts (Fig. S7). Previous work investigating the RNA binding landscape of a panel of hnRNP proteins in a non-macrophage cell line (HEK293Ts, human embryonic kidney cells) revealed that hnRNP M has a strong preference for binding distal intronic regions (>2kb from an exon-intron junction)^60^. Its binding profile was somewhat unique among the hnRNPs queried and was more reminiscent of another RNA binding protein, TDP-43. TDP-43 also binds UG-rich sites in distal introns and is crucial for repressing splicing of cryptic exons for a set of transcripts in the brain^61,62^. We speculate that hnRNP M regulates splicing of macrophages transcripts through a similar mechanism where it binds to UG-rich regions downstream of cryptic exons and inhibits assembly of the spliceosome on these introns, thus slowing intron removal.

To fully elucidate how hnRNP M represses splicing at cryptic exon-containing transcripts, future experiments will need to identify the spliceosomal protein-binding partners of hnRNP M in macrophages. To date, reports in other cell types demonstrate that hnRNP M can bind to a number of splicing factors to influence RNA processing. In HeLa cells, hnRNP M can directly interact with components of the nineteen complex (CDC5L and PLRG1)^37^, as well as the polypyrimidine tract-binding protein-associated splicing factor (PSF) and the related protein p54(nrb)^33^. Studies in mesenchymal cell lines have demonstrated a critical role for hnRNP M in co-regulating alternative splicing during the epithelial-mesenchymal transition (EMT) with another protein, ESRP1 (epithelial splicing regulatory protein 1), by competing for the same polyGU-rich motif^63^. It is likely that hnRNP M functions similarly in macrophages, either by competing with splicing activators at pre-mRNAs like *IL6*, or by repressing spliceosomal recruitment/assembly, as was recently purported for hnRNP L^64^. Curiously, the hnRNP M-dependent transcript regulon we report in RAW 264.7 macrophages bears little resemblance to those reported in other cell types^27,60^, likely reflecting a cell type-specific catalog of hnRNP M protein-binding partners that can be regulated via phosphorylation downstream of innate immune sensing.

While hnRNP M’s ability to associate with the *IL6* genomic locus via chromatin immunoprecipitation is RNA-dependent, it is conceivable that hnRNP M controls innate immune gene expression through mechanisms that are independent of direct contacts between hnRNP M and regulated transcripts. Because a number of splicing factors have been shown to impact histone markers and chromatin remodeling, it is possible that hnRNP M promotes epigenetic changes at specific target transcripts^65–68^. hnRNP M may also interact with one or more lncRNAs, a number of which are regulated by TLR activation^69^ and have been shown to control *IL6* expresion^69,70^. Experiments designed to identify hnRNP M-associated RNAs in uninfected and infected macrophages will provide important insights into how hnRNP M recognizes chromatin-associated target transcripts and help illuminate how pre-mRNA splicing decisions shape the innate immune transcriptome.

## Acknowledgements

We would like to thank Drs. Jeffery Cox, Bennett Penn, and Jonathan Budzik for generously sharing *M. tuberculosis* phosphoproteomics data as well as members of the Watson and Patrick labs for critical reading of the manuscript and invaluable feedback. We would also like to thank the Texas A&M AgriLife Genomics & Bioinformatics Services for performing RNA sequencing experiments and providing computing resources.

## Methods

### Cell Lines and Culture Conditions

RAW 264.7 macrophages (ATCC) were cultured at 37°C with a humidified atmosphere of 5% CO_2_ in DMEM (Thermo Fisher) with 10% FBS (Sigma Aldrich) 0.5% HEPES (Thermo Fisher). For RAW 264.7 macrophages stably expressing scramble knockdown and hnRNP M knockdown, cells were transfected with scramble non-targeting shRNA constructs and hnRNP M shRNA constructs targeted towards the 3’ UTR of hnRNP M. After 48 hours, media was supplemented with hygromycin (Invitrogen) to select for cells containing the shRNA plasmid. RAW 264.7 macrophages stably expressing GFP-FL and hnRNP M-FL were transfected for 48 hours and then selected through addition of puromycin (Invivogen).

### LPS Treatment

RAW 264.7 macrophages were plated on 12-well tissue-culture treated plates at a density of 7.5×10^5^ and allowed to acclimate overnight. Cells were then treated with E. Coli Lipopolysaccharide (Sigma-Aldrich) at 100ng/mL. for the respective time points where supernatants and RNA were collected for analysis.

### S. Typhimurium Infection

*Salmonella enterica serovar* Typhimurium (SL1344) was obtained from Helene Andrews-Polymenis, TAMHSC. Infections with S. Typhimurium were conducted by plating RAW 264.7 macrophages on tissue-cultured treated 12-well dishes at 7.5×10^5^ and incubated overnight. Overnight cultures of S. Typhimurium were diluted 1:20 in LB broth containing 0.3M NaCl and grown until they reached an OD600 of 0.9. Unless specified, cell lines at a confluency of 80% were infected with the S. Typhimurium strains at an MOI of 10 for 30 minutes in Hank’s buffered salt solution (HBSS), and subsequently cells were spun for 10 minutes at 1,000rpm, washed twice in HBSS containing 100μg/ml of gentamycin, and refilled with media plus gentamicin (10 μg/ml). Supernatants were collected at 2 hours and 4 hours and analyzed using IL-6 (Biolegend). After removal of supernatant, cells were lysed in Trizol (Thermo Fisher) for RNA collection and analyzed using RT-qPCR.

### Immunofluorescence Microscopy

RAW 264.7 macrophages were plated on glass coverslips in 48-well plates. Cells were treated with LPS as described above. At the designated time points, cells were washed with PBS (Thermo Fisher) and then fixed in 4% paraformaldehyde for 10 minutes. Cells were washed with PBS 3x and then permeabilized with 0.2% Triton-X (Thermo Fisher). Coverslips were placed in primary antibody for 1 hour then washed 3x in PBS and placed in secondary antibody. These were washed twice in PBS and twice in deionized water, followed by mounting onto a glass slide using ProLong Diamond antifade mountant (Invitrogen). Images were acquired on a Nikon A1-Confocal Microscope.

### Western Blots

Protein samples were run on Any kD Mini-PROTEAN TGX precast protein gels (BioRad) and transferred to .45 um nitrocellulose membranes (GE Healthcare). The membranes were incubated in the primary antibody of interest overnight and washed with TBS-Tween 20. Membranes were then incubated in secondary antibody for 1-2 hours and imaged using LI-COR Odyssey FC Imaging System.

### Antibodies

The following primary antibodies were used: rabbit polyclonal hnRNP M (Abcam, #177957), rabbit polyclonal Histone 3 (Abcam, #1791), mouse monoclonal Beta-Actin (Abcam, #6276), mouse monoclonal hnRNP L (Abcam, #6106-100), rabbit polyclonal Beta-Tubulin (Abcam, #179513), mouse monoclonal hnRNP U (Santa-Cruz, sc-32315), DAPI nuclear staining (Thermo Fisher), and mouse monoclonal ANTI-FLAG M2 antibody (Sigma-Aldrich, F3165). Secondary antibodies used were as follows: IR Dye CW 680 goat anti-rabbit, IR Dye CW800 goat anti-mouse (LI-COR), Alexfluor-488 anti-rabbit and Alexafluo-647 anti-mouse secondary antibodies for immunofluorescence (LI-COR).

### Cellular fractionation and RNA isolation

Macrophage cellular fractionation was done as described in Pandya-Jones et al., 2013. Briefly, RAW 264.7 macrophages were plated on in 10 cm tissue-culture treated plates at 1-3×10^7^ per plate. Cells and buffers were kept on ice unless noted otherwise. Cells were rinsed twice in cold PBS-EDTA (Lonza) and scraped into 15-ml conical tubes. Cells were spun at 1,000 g for 5 minutes at 4C and resuspended in NP-40 lysis buffer (10 mM Tris-HCl [pH 7.5], 0.05% NP40 [Sigma], 150 mM NaCl, protease inhibitor tablet (Thermo Fisher)) and incubated for 5 min on ice. Lysate was added to 2.5 volumes of a sucrose cushion (Lysis buffer with 24% sucrose) and centrifuged for at 14,000 rpm for 10min at 4C. The supernatant was collected and saved for cytoplasmic protein sample. The nuclear pellet was resuspended in glycerol buffer (20 mM Tris-HCl [pH 7.9], 75 mM NaCl, 0.5 mM EDTA, 0.85 mM DTT, 50% glycerol, protease inhibitor tablet) and lysed with nuclear lysate buffer in equal volume and vortexed 2X for 2 seconds (10 mM HEPES [pH 7.6], 1 mM DTT, 7.5 mM MgCl_2_, 0.2 mM EDTA, 0.3 M NaCl, 1 M UREA, 1% NP-40, protease inhibitor tablet). Lysates were chilled on ice for 2 minutes and then spun at 10,000 rpm for 2 minutes at 4C. Supernatant was collected and used for nucleoplasmic protein samples. The remining chromatin pellet was gently rinsed in PBS-EDTA and treated with DNase in DNase buffer for 1 hr at 37C. After incubation, the supernatant was collected for chromatin protein samples. Sample buffer (BIO-RAD) and 2-Mercapoethanol (BIO-RAD) was added to every protein sample with 5 minutes boiling prior to running on gels for western blots. Approximately 10% of sample was loaded for western blots.

### RNase Fractionation

For nuclear lysates treated with RNase, nuclear pellets were responded in glycerol buffer. Nuclear lysis buffer was added, and lysates were incubated on ice for 5 minutes. Samples were then divided into two samples with one receiving 1 ul of RNAse A (Thermo Fisher) per 50 ul sample and another with no RNAse A. Both were incubated at 37C for 30 mins. Lysates were then spun at 10,000rpm for 2 mins and the rest of the fractionation proceeded as described.

### Gene Ontology (GO) Canonical Pathway Analysis

To determine the most affected pathways in control vs. hnRNP M knockdown RAW 264.7 macrophages, canonical pathway analysis was conducted using Ingenuity Pathway Analysis software from Qiagen Bioinformatics. Genes that were differentially expressed wth a p-value<0.05 from our RNA-SEQ analysis were used as input from uninfected and Salmonella Typhimurium infected cells. The top hits were represented in bar graphs by z-score.

### RNA isolation and qPCR analysis

For transcript analysis, cells were harvested in Trizol and RNA was isolated using Direct-zol RNA Miniprep kits (Zymo Research) with 1 hr DNase treatment. cDNA was synthesized with iScript cDNA Synthesis Kit (Bio-Rad). CDNA was diluted to 1:20 for each sample. A pool of cDNA from each treated or infected sample was used to make a 1:10 standard curve with each standard sample diluted 1:5 to produce a linear curve. RT-qPCR was performed using Power-Up SYBR Green Master Mix (Thermo Fisher) using a Quant Studio Flex 6 (Applied Biosystems). Samples were run in triplicate wells in a 96-well plate. Averages of the raw values were normalized to average values for the same sample with the control gene, beta-actin. To analyze fold induction, the average of the treated sample was divided by the untreated control sample, which was set at 1.

### Chromatin Immunoprecipitation

Chromatin Immunoprecipitation (ChIP) was adapted from Abcam’s protocol. Briefly, two confluent 15 cm dishes of RAW 264.7 macrophages were crosslinked in formaldehyde to a final concentration of 0.75% and rotated for 10 minutes. Glycine was added to stop the cross linking by shaking for 5 minutes at a concentration of 125 mM. Cells were rinsed with PBS twice and then scraped into 5 mL PBS and centrifuged at 1,000g for 5 min at 4C. Cellular pellets were resuspended in ChIP lysis buffer (750 μL per 1×10^7^ cells) and incubated for 10 min on ice. Cellular lysates were sonicated for 40 minutes (30sec ON, 30sec OFF) on high in a Bioruptor UCD-200 (Diagenode). After sonication, cellular debris was pelleted by centrifugation for 10 min, 4°C, 8,000 × g. Input samples were taken at this step and stored at −80C until decrosslinking. For RNase treated samples, RNase A was added to cell lysates and incubated for 30 mins at 37C. Approximately 25 μg of DNA diluted to 1:10 with RIPA buffer was used for overnight immunoprecipitation. Each ChIP had one sample for the specific antibody and one sample for Protein G beads only which were pre-blocked for 1 hr with single stranded herring sperm DNA (75 ng/μL) and BSA (0.1 μg/μL). The respective primary antibody was added to all samples except the beads-only sample at a concentration of 5 ug and rotated at 4°C overnight. Beads were washed 3x in with a final wash in high salt (500mM NaCl). DNA was eluted with elution buffer and rotated for 15 min at 30C. Centrifuge for 1 min at 2,000 × g and transfer the supernatant into a fresh tube. Supernatant was incubated in NaCl, RNase A (10 mg/mL) and proteinase K (20 mg/mL) and incubated at 65°C for 1 h. The DNA was purified using phenol:chloroform extraction. DNA levels were measure by RT-qPCR. Primers were designed by tiling each respective gene every 500 base pairs that were inputted into NCBI primer design.

### FLAG Chromatin Immunoprecipitation

In RAW 264.7 macrophages stably expressing hnRNP M-FL and GFP-FL, ChIP was conducted as described above with minor adjustments. Lysates were incubated overnight at 4C with ANTIFLAG M2 antibody. After washing, DNA was eluted with FLAG peptide (Sigma-Aldrich F4799) by adding 20 ul of 5X FLAG peptide, vortexed at room temperature for 15 mins and supernatants were collected. This process was repeated a total of 3x followed by decrosslinking as described.

### RNA-SEQ

Total RNA was extracted as previously described above. Preparation and sequencing of cDNA libraries were done by Texas A&M AgriLife Genomics and Bioinformatics Service using the Illumina HiSeq 4000. Reads were mapped to the reference genome sequence of Mus musculus from GRCm38 (RefSeq) using the CLC Genomics Workbench 8.0.1 (Qiagen). Relative transcript expression was calculated by counting the Reads Per Kilobase of exon model per Million mapped reads (RPKM). Statistical Analysis was conducted comparing scramble control cells vs. hnRNP M KD cells in uninfected and S. Typhimurium infected samples through CLC Genomics Workbench Empirical Analysis of DGE. Genes with p-values<0.05 were displayed in volcano plots and heat maps using GraphPad Prism software (GraphPad, San Diego, CA).

### Alternative Splicing Analysis

Alternative splicing events were analyzed using MAJIQ and VOILA with the default parameters (Vaquero-Garcia, 2016). Briefly, uniquely mapped, junction-spanning reads were used by MAJIQ to construct splice graphs for transcripts by using the RefSeq annotation supplemented with de-novo detected junctions. Here, de-novo refers to junctions that were not in the RefSeq transcriptome database but had sufficient evidence in the RNA-Seq data. The resulting gene splice graphs were analyzed for all identified local splice variations (LSVs). For every junction in each LSV, MAJIQ then quantified expected percent spliced in (PSI) value in control and hnRNP M knockdown samples and expected change in PSI (dPSI) between control and hnRNP M KD samples. Results from VOILA were then filtered for high confidence changing LSVs (whereby one or more junctions had at least a 95% probability of expected dPSI of at least an absolute value of 20 PSI units (noted as “20% dPSI”) between control and hnRNP M KD) and candidate changing LSVs (95% probability, 10% dPSI). For the high confidence results (dPSI >= 20%), the events were further categorized as single exon cassette, multi-exon cassette, alternative 5’ and/or 3’ splice site, intron-retention.

### VSV infection

7×10^5^ RAW cells were seeded in 12-well plates 16h before infection. Cells were infected with VSV-GFP virus^71^ at multiplicity of infection (MOI) of 10, 1 and 0.1 in serum-free DMEM (HyClone SH30022.01). After 1h of incubation with media containing virus, supernatant was removed, and fresh DMEM plus 10% FBS was added to each well. At indicated times post infection, cells were harvested with Trizol and prepared for RNA isolation.

### Statistics

Statistical analysis of data was performed using GraphPad Prism software. Two-tailed unpaired Student’s t-tests were used for statistical analyses, and unless otherwise noted, all results are representative of at least three biological experiments and are reported as the mean ± SEM (n = 3 per group).

## Author Contributions

K.O.W, K.L.P., and R.O.W. designed the project. K.O.W. performed the experiments and analyzed the data. H. M. S. performed exon-inclusion PCR assays. S.T. and A.P.W. performed VSV infections. K.O.W., K.L.P., and R.O.W. wrote the paper and created figures. All authors read and approved the final version of the manuscript.

## Competing Interests

The authors declare no competing interests.

## Materials and Correspondence

Correspondence and requests for materials should be addressed to K.L.P. or R.O.W.

## Data Availability

The authors declare that the data supporting the findings of this study are available within the article and its Supplementary Information files or are available from the authors upon request.

## Notes

#### Summary of Updates

Figure 6 updated; portions of text updated/clarified

## References

1. Bhatt, D. M. et al. Transcript dynamics of proinflammatory genes revealed by sequence analysis of subcellular RNA fractions. Cell 150, 279–290 (2012).

2. Pandya-Jones, A. et al. Splicing kinetics and transcript release from the chromatin compartment limit the rate of Lipid A-induced gene expression. RNA 19, 811–827 (2013).

3. Pai, A. A. et al. Widespread Shortening of 3’ Untranslated Regions and Increased Exon Inclusion Are Evolutionarily Conserved Features of Innate Immune Responses to Infection. PLoS Genet. 12, e1006338 (2016).

4. Janssens, S., Burns, K., Vercammen, E., Tschopp, J. & Beyaert, R. MyD88S, a splice variant of MyD88, differentially modulates NF-kappaB-and AP-1-dependent gene expression. FEBS Lett. 548, 103–107 (2003).

5. Rao, N., Nguyen, S., Ngo, K. & Fung-Leung, W.-P. A novel splice variant of interleukin-1 receptor (IL-1R)-associated kinase 1 plays a negative regulatory role in Toll/IL-1R-induced inflammatory signaling. Mol. Cell. Biol. 25, 6521–6532 (2005).

6. Seo, J.-W., Yang, E.-J., Kim, S. H. & Choi, I.-H. An inhibitory alternative splice isoform of Toll-like receptor 3 is induced by type I interferons in human astrocyte cell lines. BMB Rep 48, 696–701 (2015).

7. Gray, P. et al. Identification of a novel human MD-2 splice variant that negatively regulates Lipopolysaccharide-induced TLR4 signaling. J. Immunol. 184, 6359–6366 (2010).

8. De Arras, L. & Alper, S. Limiting of the innate immune response by SF3A-dependent control of MyD88 alternative mRNA splicing. PLoS Genet. 9, e1003855 (2013).

9. De Arras, L. et al. An evolutionarily conserved innate immunity protein interaction network. J. Biol. Chem. 288, 1967–1978 (2013).

10. Zhao, W. et al. Nuclear to cytoplasmic translocation of heterogeneous nuclear ribonucleoprotein U enhances TLR-induced proinflammatory cytokine production by stabilizing mRNAs in macrophages. J. Immunol. 188, 3179–3187 (2012).

11. Chen, Y.-L. et al. Transcriptional regulation of tristetraprolin by NF-κB signaling in LPS-stimulated macrophages. Mol. B¡oi Rep. 40, 2867–2877 (2013).

12. Ostareck, D. H. & Ostareck-Lederer, A. RNA-Binding Proteins in the Control of LPS-Induced Macrophage Response. Front Genet 10, 31 (2019).

13. Liepelt, A. et al. Translation control of TAK1 mRNA by hnRNP K modulates LPS-induced macrophage activation. RNA 20, 899–911 (2014).

14. Allemand, E. et al. Regulation of heterogenous nuclear ribonucleoprotein A1 transport by phosphorylation in cells stressed by osmotic shock. Proc. Natl. Acad. Sci. U.S.A. 102, 3605–3610 (2005).

15. Shin, C., Feng, Y. & Manley, J. L. Dephosphorylated SRp38 acts as a splicing repressor in response to heat shock. Nature 427, 553–558 (2004).

16. Huang, Y., Yario, T. A. & Steitz, J. A. A molecular link between SR protein dephosphorylation and mRNA export. Proc. Natl. Acad. Sci U.S.A. 101, 9666–9670 (2004).

17. Cobianchi, F., Calvio, C., Stoppini, M., Buvoli, M. & Riva, S. Phosphorylation of human hnRNP protein A1 abrogates in vitro strand annealing activity. Nucleic Acids Res. 21, 949–955 (1993).

18. Ostareck-Lederer, A. et al. c-Src-mediated phosphorylation of hnRNP K drives translational activation of specifically silenced mRNAs. Mol. Cell. Biol. 22, 4535–4543 (2002).

19. Stamm, S. Regulation of alternative splicing by reversible protein phosphorylation. J. Biol. Chem. 283, 1223–1227 (2008).

20. Penn, B. H. et al. An Mtb-Human Protein-Protein Interaction Map Identifies a Switch between Host Antiviral and Antibacterial Responses. Mol. Cell 71, 637–648.e5 (2018).

21. Pandey, A. et al. Global Reprogramming of Host Kinase Signaling in Response to Fungal Infection. Cell Host Microbe 21, 637–649.e6 (2017).

22. Passacantilli, I., Frisone, P., De Paola, E., Fidaleo, M. & Paronetto, M. P. hnRNPM guides an alternative splicing program in response to inhibition of the PI3K/AKT/mTOR pathway in Ewing sarcoma cells. Nucleic Acids Res. 45, 12270–12284 (2017).

23. Thomas, P., Forse, R. A. & Bajenova, O. Carcinoembryonic antigen (CEA) and its receptor hnRNP M are mediators of metastasis and the inflammatory response in the liver. Clin. Exp. Metastasis 28, 923–932 (2011).

24. Chen, S. et al. Identification of HnRNP M as a novel biomarker for colorectal carcinoma by quantitative proteomics. Am. J. Physiol Gastrointest. Liver Physiol 306, G394–403 (2014).

25. Xu, Y. et al. Cell type-restricted activity of hnRNPM promotes breast cancer metastasis via regulating alternative splicing. Genes Dev. 28, 1191–1203 (2014).

26. Chen, W.-Y. et al. Heterogeneous nuclear ribonucleoprotein M associates with mTORC2 and regulates muscle differentiation. Sci Rep 7, 41159 (2017).

27. Damianov, A. et al. Rbfox Proteins Regulate Splicing as Part of a Large Multiprotein Complex LASR. Cell 165, 606–619 (2016).

28. Jagdeo, J. M. et al. Heterogeneous Nuclear Ribonucleoprotein M Facilitates Enterovirus Infection. J. Virol 89, 7064–7078 (2015).

29. LaPointe, A. T., Gebhart, N. N., Meller, M. E., Hardy, R. W. & Sokoloski, K. J. Identification and Characterization of Sindbis Virus RNA-Host Protein Interactions. J. Virol 92, 1137 (2018).

30. Viktorovskaya, O. V., Greco, T. M., Cristea, I. M. & Thompson, S. R. Identification of RNA Binding Proteins Associated with Dengue Virus RNA in Infected Cells Reveals Temporally Distinct Host Factor Requirements. PLoS Negl Trop Dis 10, e0004921 (2016).

31. Yamamoto, M. et al. Role of adaptor TRIF in the MyD88-independent toll-like receptor signaling pathway. Science 301, 640–643 (2003).

32. Hovhannisyan, R. H. & Carstens, R. P. Heterogeneous ribonucleoprotein m is a splicing regulatory protein that can enhance or silence splicing of alternatively spliced exons. J. Biol. Chem. 282, 36265–36274 (2007).

33. Marko, M., Leichter, M., Patrinou-Georgoula, M. & Guialis, A. hnRNP M interacts with PSF and p54(nrb) and co-localizes within defined nuclear structures. Exp. Cell Res. 316, 390–400 (2010).

34. Sun, H., Kamanova, J., Lara-Tejero, M. & Galán, J. E. Salmonella stimulates pro-inflammatory signalling through p21-activated kinases bypassing innate immune receptors. Nat Microbiol 3, 1122–1130 (2018).

35. Poltorak, A. et al. Defective LPS signaling in C3H/HeJ and C57BL/10ScCr mice: mutations in Tlr4 gene. Science 282, 2085–2088 (1998).

36. Stetson, D. B. & Medzhitov, R. Recognition of cytosolic DNA activates an IRF3-dependent innate immune response. Immunity 24, 93–103 (2006).

37. Llères, D., Denegri, M., Biggiogera, M., Ajuh, P. & Lamond, A. I. Direct interaction between hnRNP-M and CDC5L/PLRG1 proteins affects alternative splice site choice. EMBO Rep. 11, 445–451 (2010).

38. Park, E. et al. Regulatory roles of heterogeneous nuclear ribonucleoprotein M and Nova-1 protein in alternative splicing of dopamine D2 receptor pre-mRNA. J. Biol. Chem. 286, 25301–25308 (2011).

39. Cho, S. et al. hnRNP M facilitates exon 7 inclusion of SMN2 pre-mRNA in spinal muscular atrophy by targeting an enhancer on exon 7. Biochim. Biophys. Acta 1839, 306–315 (2014).

40. Vaquero-Garcia, J. et al. A new view of transcriptome complexity and regulation through the lens of local splicing variations. Elife 5, e11752 (2016).

41. Pettit Kneller, E. L., Connor, J. H. & Lyles, D. S. hnRNPs Relocalize to the cytoplasm following infection with vesicular stomatitis virus. J. Virol. 83, 770–780 (2009).

42. Lichtenstein, M., Guo, W. & Tartakoff, A. M. Control of nuclear export of hnRNP A1. Traffic 2, 261–267 (2001).

43. Kosugi, S., Hasebe, M., Tomita, M. & Yanagawa, H. Systematic identification of cell cycle-dependent yeast nucleocytoplasmic shuttling proteins by prediction of composite motifs. Proc. Natl. Acad. Sci. U.S.A. 106, 10171–10176 (2009).

44. Yachdav, G. et al. PredictProtein--an open resource for online prediction of protein structural and functional features. Nucleic Acids Res. 42, W337–43 (2014).

45. Nojima, T., Gomes, T., Carmo-Fonseca, M. & Proudfoot, N. J. Mammalian NET-seq analysis defines nascent RNA profiles and associated RNA processing genome-wide. Nat Protoc 11, 413–428 (2016).

46. Pandya-Jones, A. & Black, D. L. Co-transcriptional splicing of constitutive and alternative exons. RNA 15, 1896–1908 (2009).

47. Moehle, E. A., Braberg, H., Krogan, N. J. & Guthrie, C. Adventures in time and space: splicing efficiency and RNA polymerase II elongation rate. RNA Biol 11, 313–319 (2014).

48. Moehle, E. A., Ryan, C. J., Krogan, N. J., Kress, T. L. & Guthrie, C. The yeast SR-like protein Npl3 links chromatin modification to mRNA processing. PLoS Genet. 8, e1003101 (2012).

49. Patrick, K. L. et al. Genetic interaction mapping reveals a role for the SWI/SNF nucleosome remodeler in spliceosome activation in fission yeast. PLoS Genet. 11, e1005074 (2015).

50. Nissen, K. E. et al. The histone variant H2A.Z promotes splicing of weak introns. Genes Dev. 31, 688–701 (2017).

51. Neves, L. T. et al. The histone variant H2A.Z promotes efficient cotranscriptional splicing in S. cerevisiae. Genes Dev. 31, 702–717 (2017).

52. Bieberstein, N. I., Straube, K. & Neugebauer, K. M. Chromatin immunoprecipitation approaches to determine co-transcriptional nature of splicing. Methods Mol. Biol. 1126, 315–323 (2014).

53. Kfir, N. et al. SF3B1 association with chromatin determines splicing outcomes. Cell Rep 11, 618–629 (2015).

54. Costa-Pereira, A. P. Regulation of IL-6-type cytokine responses by MAPKs. Biochem. Soc. Trans. 42, 59–62 (2014).

55. Kandasamy, R. K. et al. A time-resolved molecular map of the macrophage response to VSV infection. NPJ Syst Biol Appl 2, 16027 (2016).

56. Shin, D.-L., Hatesuer, B., Bergmann, S., Nedelko, T. & Schughart, K. Protection from Severe Influenza Virus Infections in Mice Carrying the Mx1 Influenza Virus Resistance Gene Strongly Depends on Genetic Background. J. Virol. 89, 9998–10009 (2015).

57. Ramirez-Carrozzi, V. R. et al. A unifying model for the selective regulation of inducible transcription by CpG islands and nucleosome remodeling. Cell 138, 114–128 (2009).

58. Masuda, K. et al. Arid5a controls IL-6 mRNA stability, which contributes to elevation of IL-6 level in vivo. Proc. Natl. Acad. Sci. U.S.A. 110, 9409–9414 (2013).

59. Higa, M. et al. Regulation of inflammatory responses by dynamic subcellular localization of RNA-binding protein Arid5a. Proc. Natl. Acad. Sci. U.S.A. 115, E1214–E1220 (2018).

60. Huelga, S. C. et al. Integrative genome-wide analysis reveals cooperative regulation of alternative splicing by hnRNP proteins. Cell Rep 1, 167–178 (2012).

61. Ling, J. P., Pletnikova, O., Troncoso, J. C. & Wong, P. C. TDP-43 repression of nonconserved cryptic exons is compromised in ALS-FTD. Science 349, 650–655 (2015).

62. Ling, J. P. et al. PTBP1 and PTBP2 Repress Nonconserved Cryptic Exons. Cell Rep 17, 104–113 (2016).

63. Harvey, S. E. et al. Coregulation of alternative splicing by hnRNPM and ESRP1 during EMT. RNA 24, 1326–1338 (2018).

64. McClory, S. P., Lynch, K. W. & Ling, J. P. HnRNP L represses cryptic exons. RNA 24, 761–768 (2018).

65. de Almeida, S. F. et al. Splicing enhances recruitment of methyltransferase HYPB/Setd2 and methylation of histone H3 Lys36. Nat. Struct. Mol. Biol. 18, 977–983 (2011).

66. Kim, S., Kim, H., Fong, N., Erickson, B. & Bentley, D. L. Pre-mRNA splicing is a determinant of histone H3K36 methylation. Proc. Natl. Acad. Sci. U.S.A. 108, 13564–13569 (2011).

67. Martins, S. B. et al. Spliceosome assembly is coupled to RNA polymerase II dynamics at the 3’ end of human genes. Nat. Struct. Mol. Biol. 18, 1115–1123 (2011).

68. Saldi, T., Cortazar, M. A., Sheridan, R. M. & Bentley, D. L. Coupling of RNA Polymerase II Transcription Elongation with Pre-mRNA Splicing. J. Mol. Biol. 428, 2623–2635 (2016).

69. Carpenter, S. et al. A long noncoding RNA mediates both activation and repression of immune response genes. Science 341, 789–792 (2013).

70. Atianand, M. K. et al. A Long Noncoding RNA lincRNA-EPS Acts as a Transcriptional Brake to Restrain Inflammation. Cell 165, 1672–1685 (2016).

71. Dalton, K. P. & Rose, J. K. Vesicular stomatitis virus glycoprotein containing the entire green fluorescent protein on its cytoplasmic domain is incorporated efficiently into virus particles. Virology 279, 414–421 (2001).

