## Supplemental Figures for "The splicing factor hnRNP M is a critical regulator of innate immune gene expression in macrophages"

**Figure S1:**

(a) Venn diagram to represent overlap between hnRNP M-regulated transcripts in uninfected and *Salmonella*-infected macrophages.

(b) Complete ingenuity pathway analysis for SCR vs. hnRNP M KD uninfected macrophages

(c) Complete ingenuity pathway analysis for SCR vs. hnRNP M KD *Salmonella*-infected macrophages

(d) Additional validation of RNA-seq results. Gene expression by RT-qPCR in SCR, hnRNP M KD 1, and hnRNP M KD 2 2h post-*Salmonella* infection for *Adora2a, Marcks, Mx1, Gbp5*, and 4h for *Adora2a* and *Marcks*.

**Figure S2:**

(a) Additional gene expression by RT-qPCR in LPS-treated cells, SCR, hnRNP M KD 1, and hnRNP M KD 2h post-treatment. *Tnfα, Marcks*, and *Gbp5*.

(b) *IL6* expression by RT-qPCR in SCR, hnRNP M KD1, and hnRNP M KD 2 8h post-LPS treatment.

**Figure S3:**

(a) Venn diagram to represent overlap between hnRNP M-regulated transcripts identified via RNA-SEQ analysis (unique genes identified in *Salmonella* and uninfected analyses) and hnRNP M-dependent LSVs identified via MAJIQ analysis (total number of unique LSV events from both *Salmonella* and uninfected conditions).

(b) Full VOILA-generated tracks for *Commd8* and *Nmt2*. Significant LSVs are shown in color.

**Figure S4:**

(a) Immunofluorescence microscopy of 3xFLAG-hnRNP M in untreated and LPS-treated macrophages (1h and 2h post-treatment)

(b) Immunofluorescence microscopy of hnRNP U in untreated and LPS-treated macrophages (2h post-treatment)

(c) Western blot of whole cell lysate, cytoplasm, nucleoplasm, and chromatin of fractionated stable 3xFL-hnRNP M-expressing macrophages over a time-course of LPS treatment.

(d) RNA sequence of *IL6* introns 2 and 3. Consensus or near-consensus hnRNP M binding sites are highlighted in yellow.

**Figure S5:**

(a) Western blot analysis of hnRNP M phoshomutants stably expressed in RAW 264.7 macrophages

(b) *IL6* expression by RT-qPCR, 2h post-LPS treatment in 3xFL-hnRNP M WT and 3xFL-hnRNP M S587A/D-expressing macrophages

(c) *Mx1* expression, 2h post-LPS treatment in 3xFL-hnRNP M WT and 3xFL-hnRNP M S587A/D-expressing macrophages

(d) Expression by RT-qPCR of *Rnf26, Slc6a4, Rnf128* for 3xFL-hnRNP M WT, hnRNP M KD 1 and 2, and 3xFL-S431A/D, S574A/D, S85A/D, S480A/D-expressing cells

(e) Western blot of whole cell lysate and chromatin of fractionated stable 3xFL-hnRNP M WT, 3xFL-hnRNP M 431A/D and 3xFL-hnRNP M 574A/D-expressing cell lines.

**Figure S6:**

(a) RT-qPCR of *Ifn* $\beta$ mRNA levels in SCR control and hnRNP M KD cells at 2h and 4h post-infection, MOI=0.1. (b) RT-qPCR of *Mx1* transcript in VSV infected SCR control and hnRNP M KD cells at 2h, 4h, and 8h post-infection, MOI=0.1. (c) RT-qPCR of *IL6* transcript in SCR control and hnRNP M KD cells at 2h, 4h, and 8h post-infection MOI=0.1. All figures are representative of 2 biological replicates.

**Figure S7:**

(a) Screenshots of IGV viewer of *Salmonella*-infected SCR- and hnRNP M KD RNA-SEQ reads at *Fcgr3, Cd276, Nfkbiz, Tnfaip3, Mir6989*. Red arrows indicate potential cryptic exons.

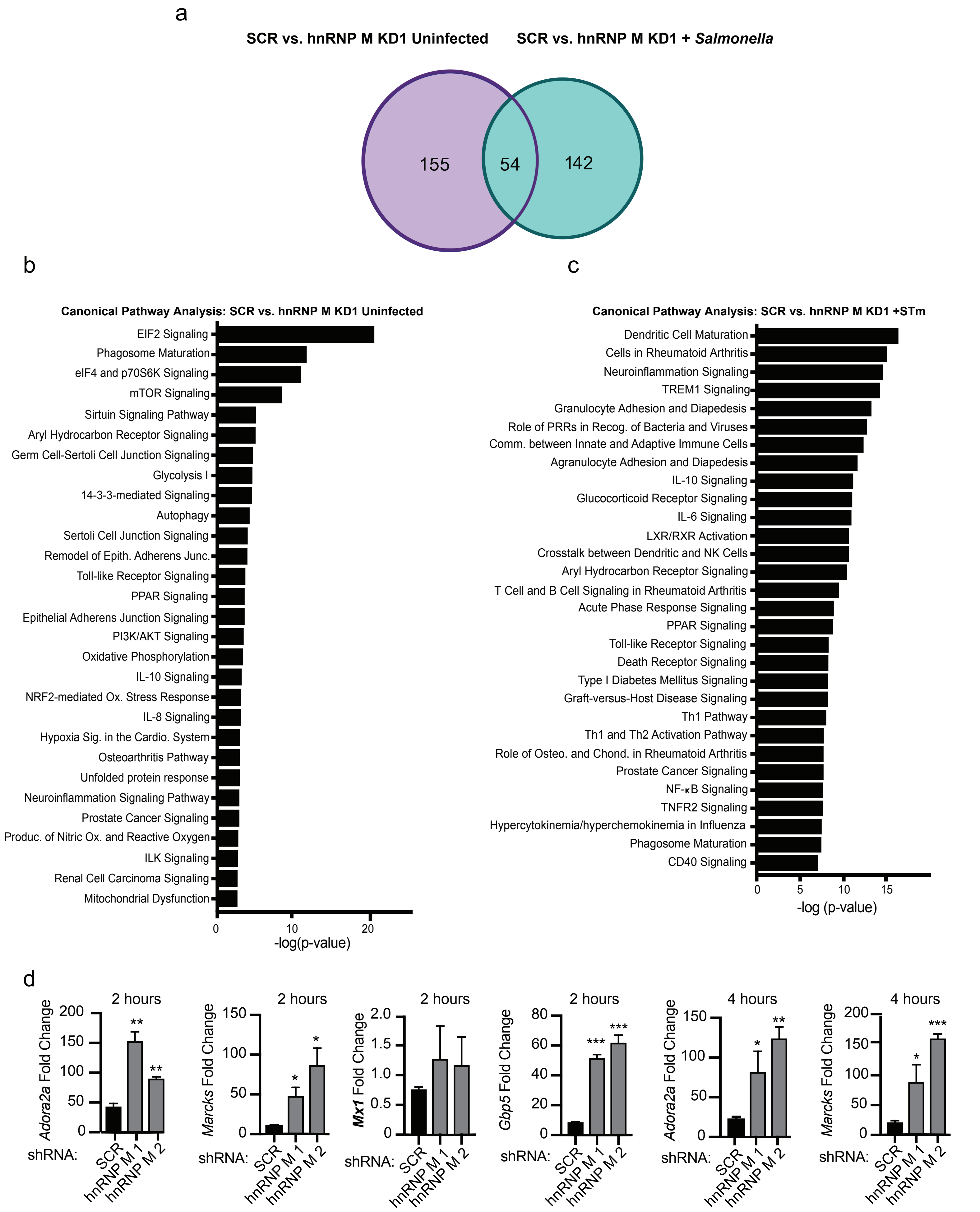


Supplemental Figure S1

a

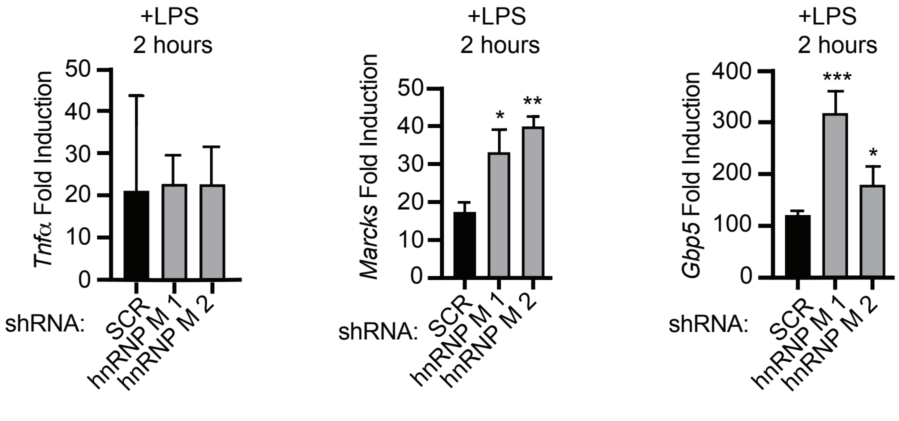


b

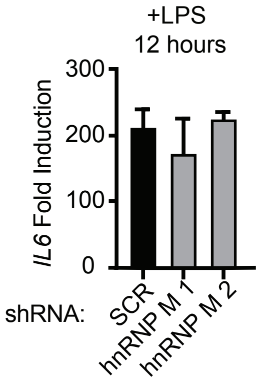


Supplemental Figure S2

a

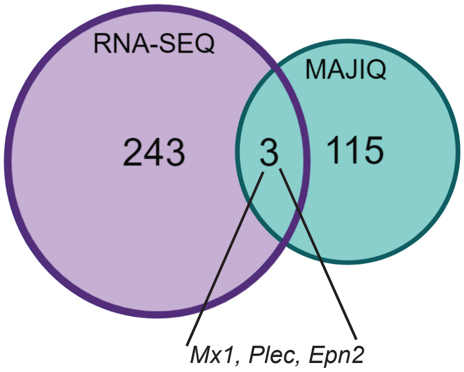


b

**Gene ID: *Commd8***

SCR

hnRNP M KD

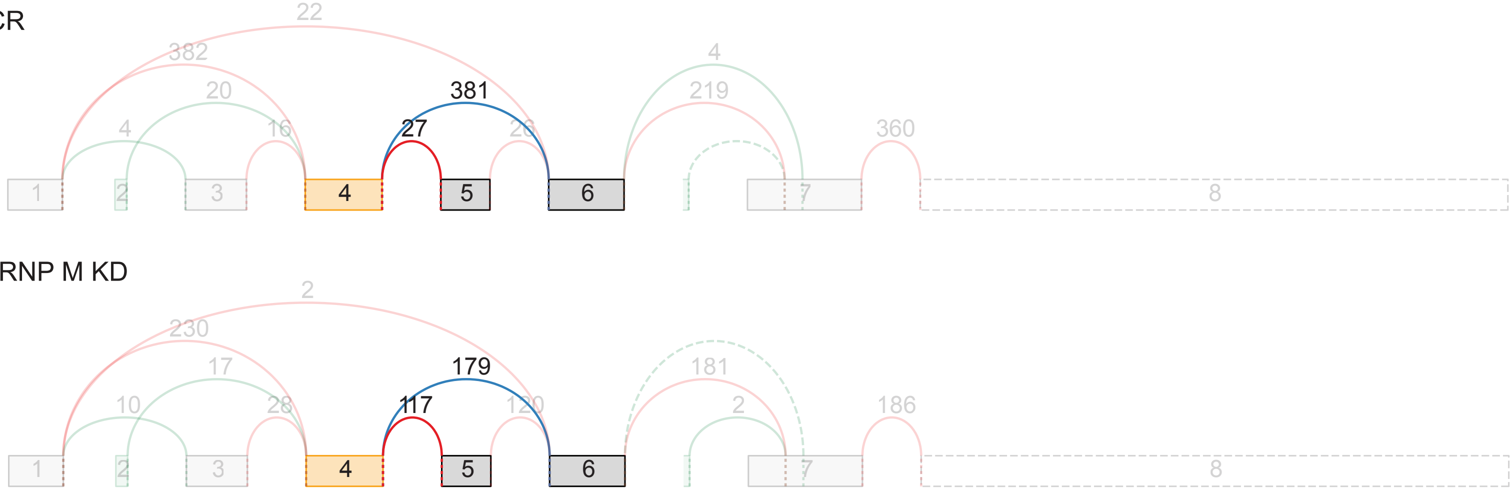


**Gene ID: *Nmt2***

SCR

hnRNP M KD

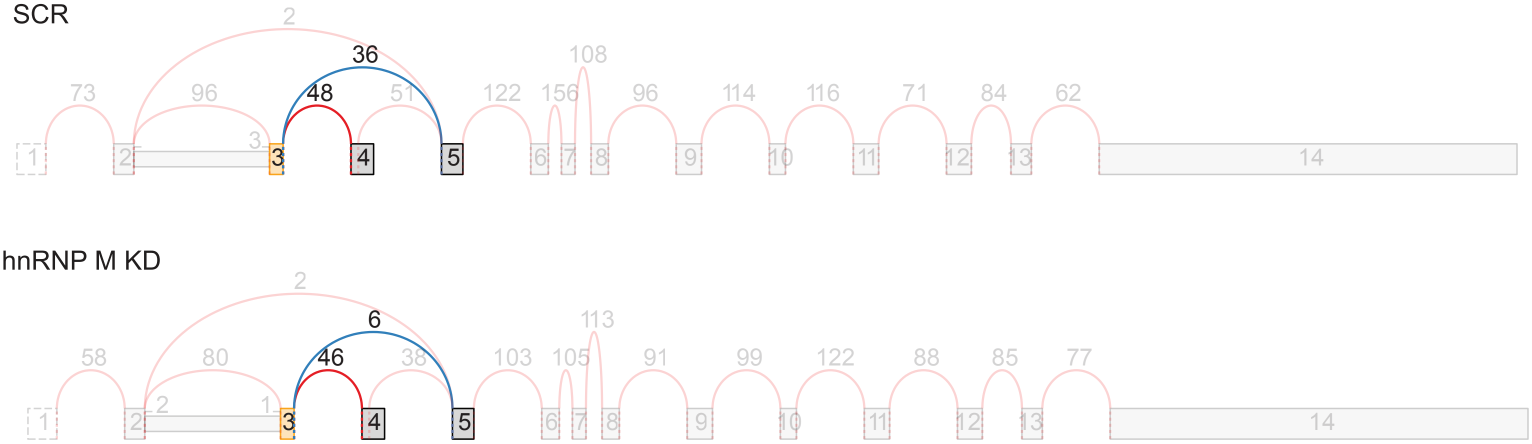


Supplemental Figure S3

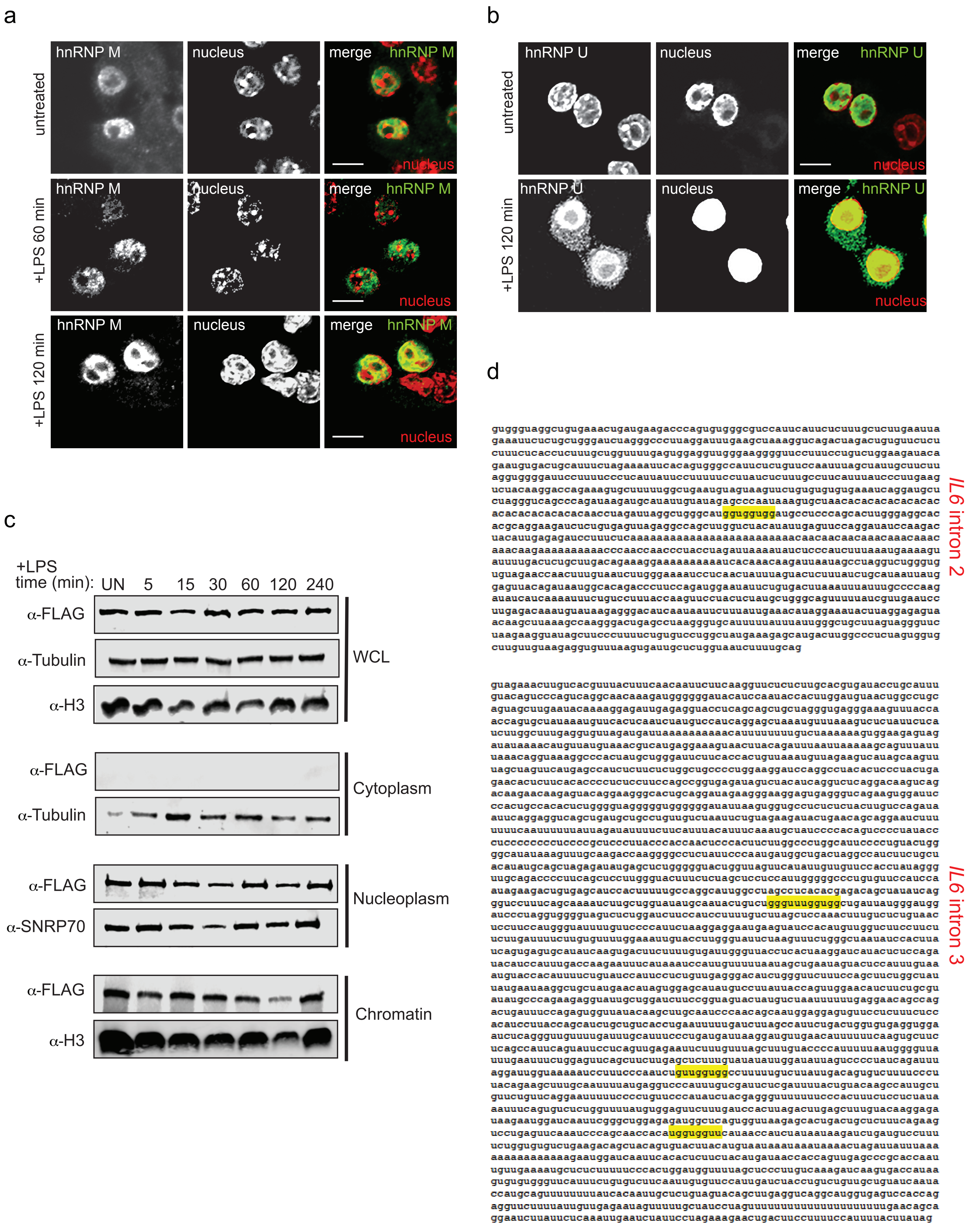


Supplemental Figure S4

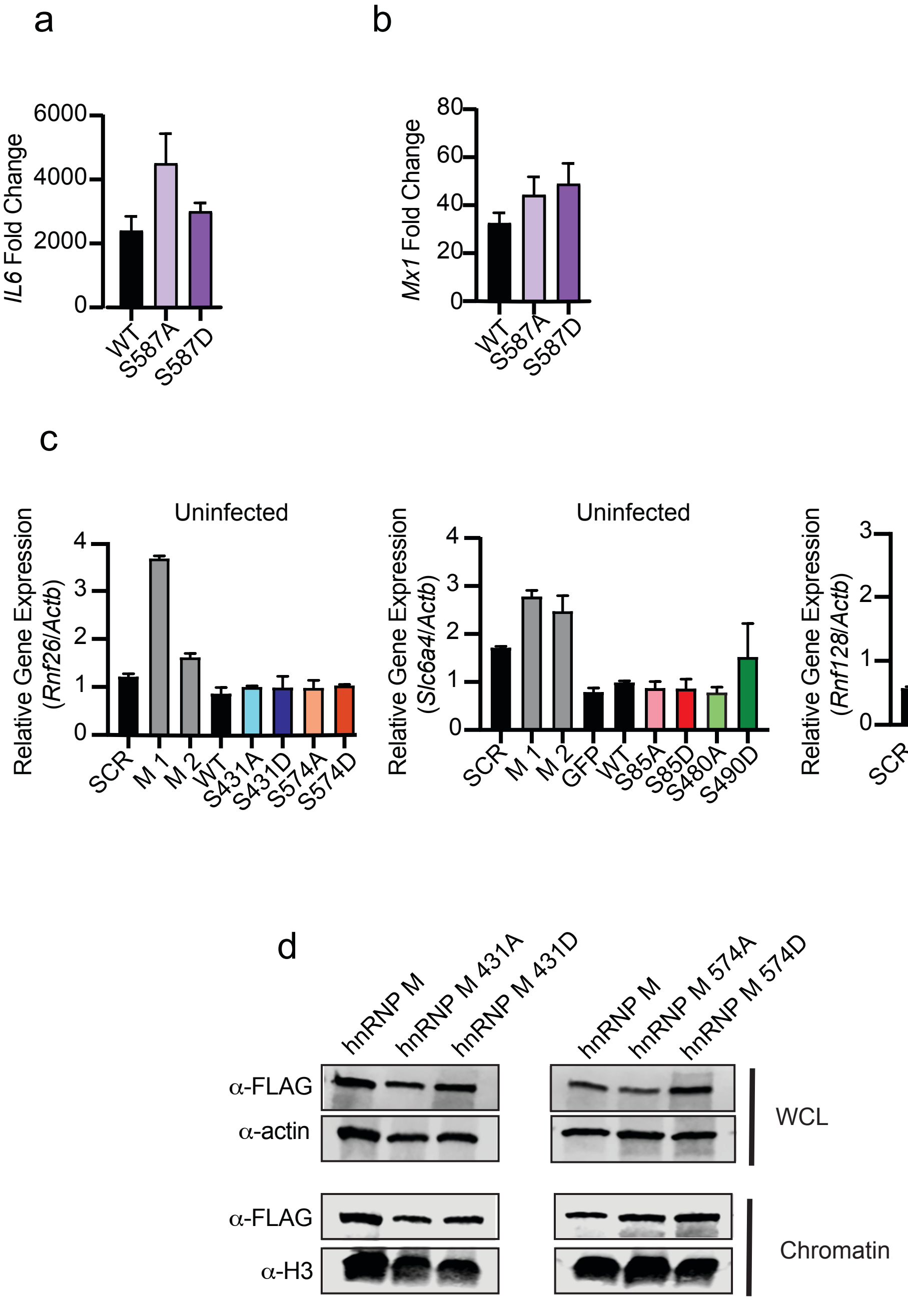


Supplemental Figure S5

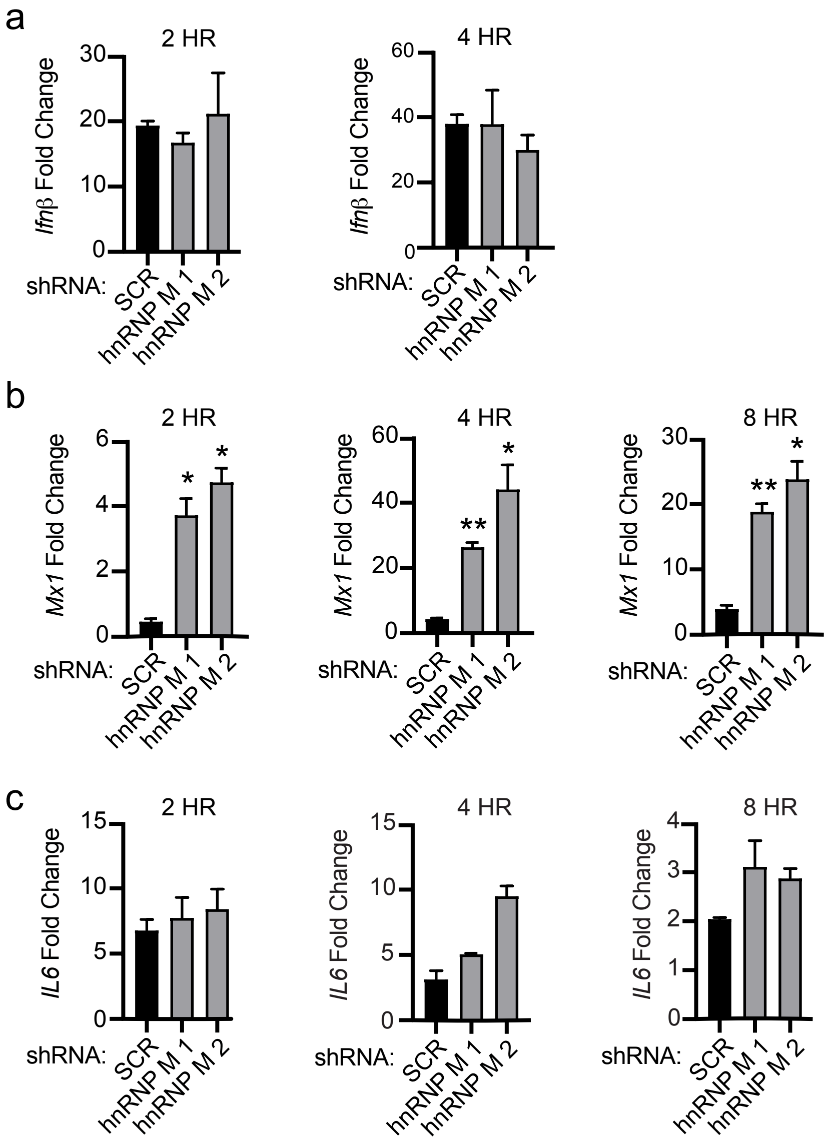


Supplemental Figure S6

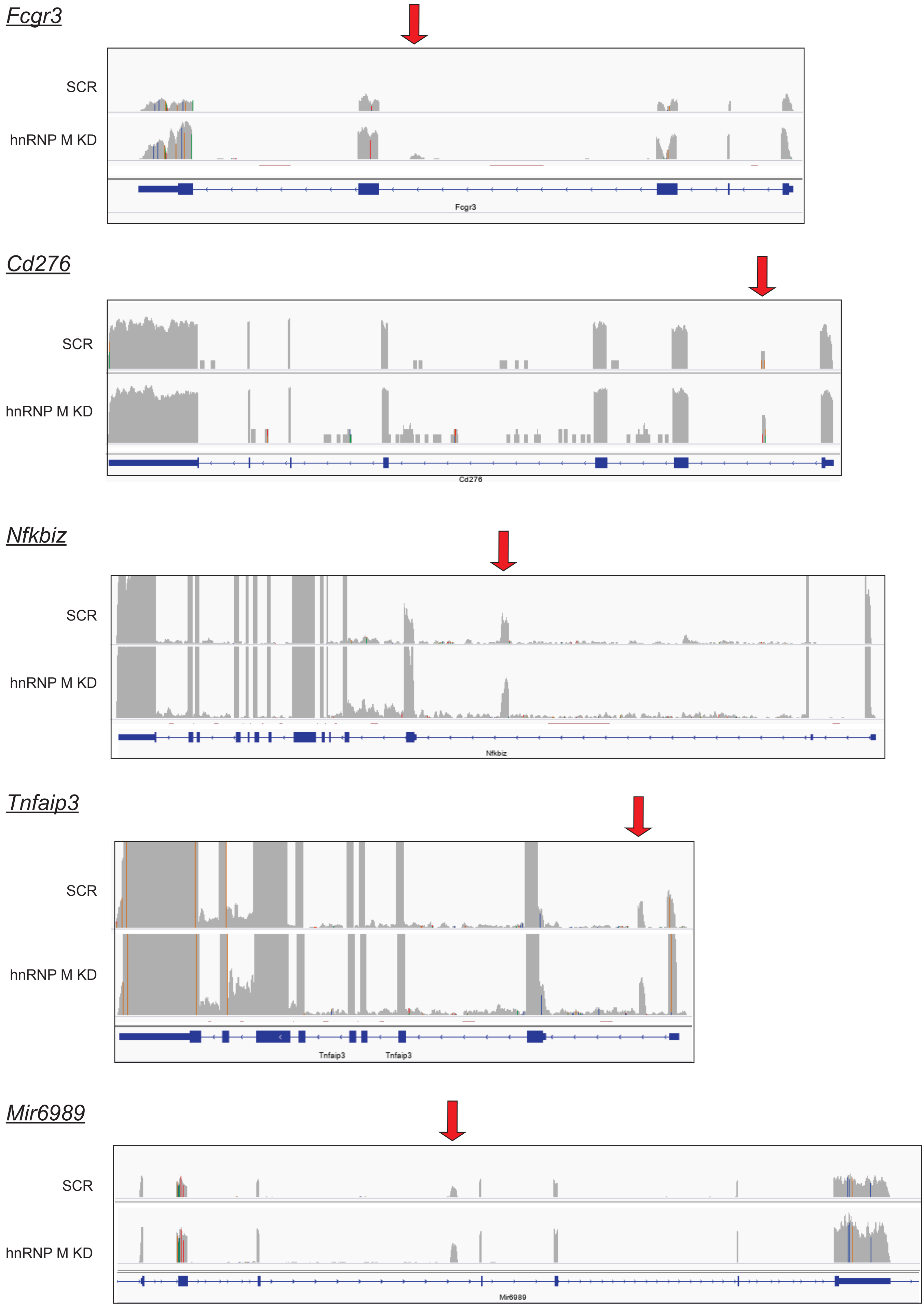


Supplemental Figure S7
